# Electro-Steric Mechanism of CLC-2 Chloride Channel Activation

**DOI:** 10.1101/2020.06.12.148783

**Authors:** José J. De Jesús-Pérez, G. Arlette Méndez-Maldonado, Irma L. Gonzalez-Hernandez, Victor De la Rosa, Roberto Gastélum-Garibaldi, Jorge E. Sánchez-Rodríguez, Jorge Arreola

## Abstract

Two-pore voltage-gated CLC chloride channels control neuronal and muscle excitability. They share a dimeric structure but their activation mechanism remains unresolved. Here we determine the step-by-step activation mechanism of the broadly expressed CLC-2 channel using homology modelling, molecular dynamic simulations and functional studies. We establish that a two-leaf gate formed by Tyr561-H_2_O-Glu213 flanked by Lys568/Glu174 and Lys212 closes the canonical pore. Activation begins when a hyperpolarization-propelled intracellular chloride occupies the pore and splits Tyr561-H_2_O-Glu213 by electrostatic/steric repulsion. Unrestrained Glu213 rotates outwardly to bind Lys212 but the pore remains closed. Protonation breaks the Glu213-Lys212 interaction while another chloride occupies the pore thus catalysing chloride exit via Lys212. Also, we found that the canonical pore is uncoupled from a cytosolic cavity by a Tyr561-containing hydrophobic gate that prevents Glu213 protonation by intracellular protons. Our data provide atomistic details about CLC-2 activation but this mechanism might be common to other CLC channels.

## Introduction

Gating is a fundamental property whereby membrane channels open a permeation pathway so ions can passively flow in or out of the cells, thus ensuing electric communication(Bezanilla, 2008; Hille et al., 1999; Jiang et al., 2003; Long et al., 2005). In some channels, a sudden change in the membrane electrical field, ligand binding or lipid bilayer forces triggers a cascade of structural rearrangements that ultimately open the permeation pathway(Bezanilla, 2008, 2002; Cox et al., 2016; Goldschen-Ohm and Chanda, 2017; Hille et al., 1999; Jiang et al., 2003; Long et al., 2005). CLC chloride (Cl^−^) channels are voltage-activated or type III(Goldschen-Ohm and Chanda, 2017) channels despite lacking a canonical voltage sensor(Jentsch and Pusch, 2018), however, the molecular mechanism of voltage-dependent activation remains elusive.

The CLC protein family includes Cl^−^ channels (CLC-0, CLC-1, CLC-2, CLC-Ka, and CLC-Kb) and 2:1 Cl^−^/H^+^ exchangers that are structurally conserved(Jentsch and Pusch, 2018). They share a double pore architecture, each pore is closed by a highly conserved Glu residue called Glu_gate_ that protrudes into the permeation pathway(Dutzler et al., 2003, 2002; Park et al., 2017; Park and MacKinnon, 2018). This structure of the pore suggested that voltage gating may occur unconventionally(Schewe et al., 2016). Glu_gate_ would be open by repulsion when voltage drives a permeant anion into the pore(Chen, 2003; De Jesús-Pérez et al., 2016; Pusch et al., 1995; Sánchez-Rodríguez et al., 2012, 2010) and/or by protonation when voltage drives a proton into the pore(Niemeyer et al., 2009a; Traverso et al., 2006a). Data collected varying the extracellular anion concentration or the anion mole fraction supported the idea that CLC-0 and CLC-1 channels were gated by permeant anions repelling the Glu_gate_(Chen, 2003; Pusch et al., 1995; Rychkov et al., 1998, 1996). However, data collected under acidic cytosolic conditions led to an alternative gating mechanism in CLC-0. This alternative mechanism invokes voltage-dependent protonation of Glu_gate_ by intracellular protons(Miller, 2006; Traverso et al., 2006a), a reaction catalysed by extracellular Cl^−^ anions(Miller, 2006). Although this hypothesis may explain CLC-0 and CLC-1 gating, the gating of CLC-2 diverges from this. CLC-2 activation depends on hyperpolarization and intracellular Cl^−^ while depolarization causes deactivation(De Santiago et al., 2005; Sánchez-Rodríguez et al., 2010; Thiemann et al., 1992). Furthermore, anions unable to permeate the pore when applied from the intracellular side still support CLC-2 voltage gating(De Jesús-Pérez et al., 2016). Notable, intracellular acidification has little effect on gating. In contrast, extracellular acidification increases the apparent open probability(Niemeyer et al., 2009b) in a voltage-independent manner(Sánchez-Rodríguez et al., 2012) but CLC-2 is still activated under unfavourable protonation condition ([H^+^]_e_ = 10^−10^ M)(De Jesús-Pérez et al., 2016). Thus, protonation cannot explain voltage-dependent activation. Instead, we proposed that hyperpolarization drives Cl^−^ into the pore causing Glu_gate_ to be open by electrostatic and steric repulsion(De Jesús-Pérez et al., 2016; Sánchez-Rodríguez et al., 2010) and then CLC-2 Glu_gate_ protonation stabilizes its open conformation(Sánchez-Rodríguez et al., 2012).

Establishing an activation mechanism for CLC Cl^−^ channels has been hindered by limited structural information on these channels and the challenge that impose gathering information about a process that occurs in nanosecond time scale within a narrow pore. Here we combined quantitative functional analysis with molecular dynamics (MD) simulations to overcome these limitations and to establish the molecular mechanism of CLC-2 activation, a Cl^−^ channel highly expressed throughout the central nervous system that controls neuronal excitability and aldosterone secretion(Boccaccio et al., 2018; Jentsch and Pusch, 2018; Thiemann et al., 1992).

## Results

### A water-mediated Tyr561-Glu_gate_ ionic interaction holds the CLC-2 pores closed

To study CLC-2 activation, CLC-2^CLC-K^ and CLC-2^CLC-1^ homology models of the CLC-2 structure were built using the CLC-K (5TQQ)(Park et al., 2017) and hCLC-1 (6COY)(Park and MacKinnon, 2018) Cl^−^ channel structures as templates. Both CLC-2^CLC-K^ and CLC-2^CLC-1^ are dimers with a transmembrane domain (TMD) comprising 17 α-helices termed B-R (Fig. 1A). Helix A (residues 1 to 96), which is not in the template structures, next to the N-terminus could not be modelled. Only the CLC-2^CLC-K^ structure included the C-terminus with two cystathionine-β-synthase (CBS) domains. Both models show the classical rhombus shape of the CLC protein structures when viewed from the top. The root mean square deviation between TMD backbones of these models is 1.0 Å and becomes 3.0 Å after adding the C-terminus.

**Fig. 1.**
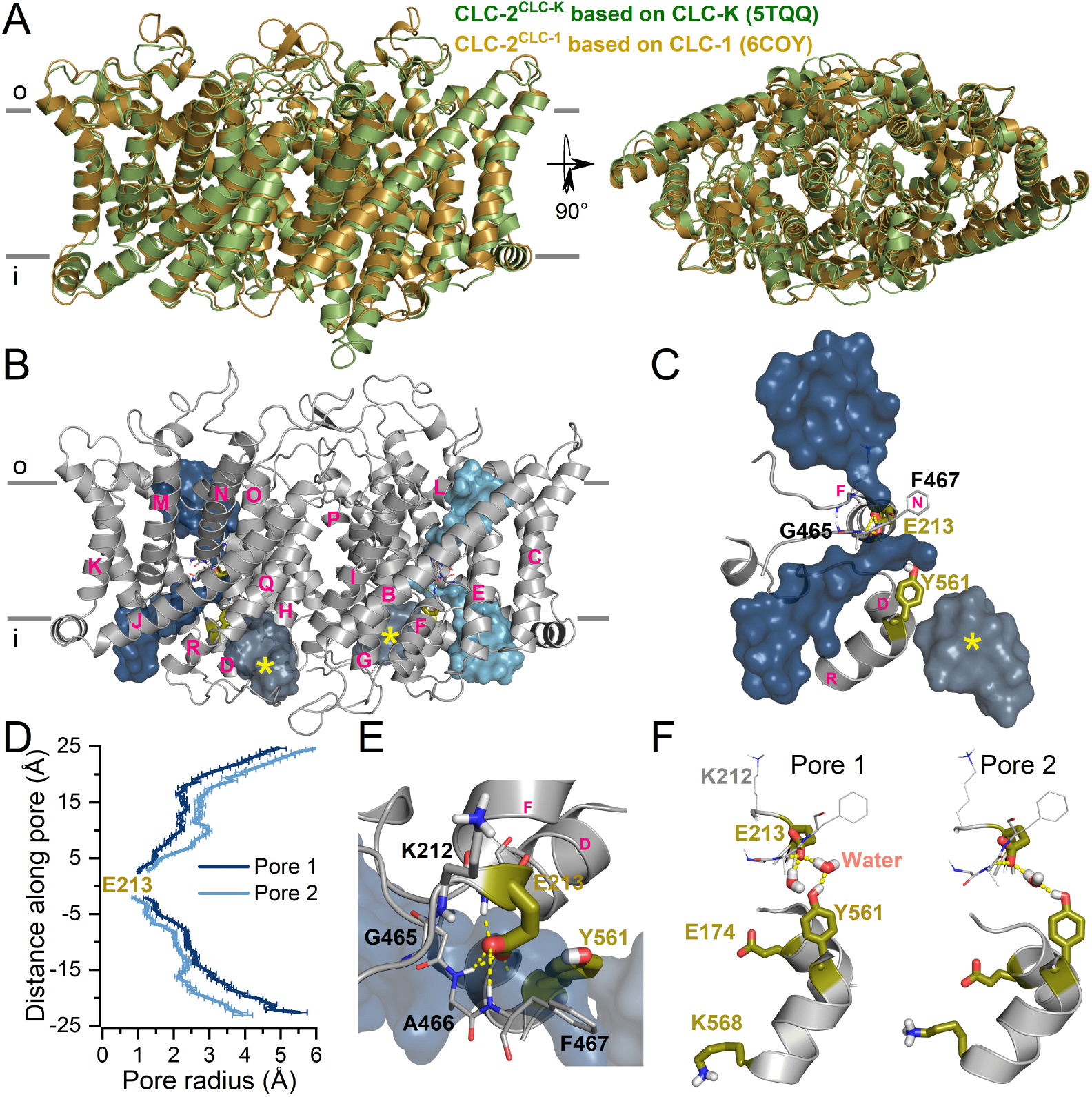
Homology models of the CLC-2 structure. A. Homodimeric models based on the cryo-EM structure of CLC-K (5TQQ, green) and hCLC-1 (6COY, orange) channels showing a view of the transmembrane domains from the membrane plane (left) or the top (right). Parallel grey lines indicate external (o) and internal (i) membrane limits. Both structures are similar (RMSD = 1.0 Å). For clarity, the intracellular carboxyl tail containing the CBS domains was not included. B. CLC-2 homology model based on the hCLC-1 structure after 30 ns of MD simulation. The names of the transmembrane helixes are shown in pink letters (B-R). The canonical pores flooded with water are represented as dark and light blue surfaces in each subunit. The intracellular alternative pathways are shown in grey marked with yellow asterisks. C. The canonical pore (dark blue) and the intracellular alternative pathway (yellow asterisk) within one subunit of CLC-2. Water molecules filling these cavities represent the structure. Tyr561 and Glu213 (Glu_gate_) are shown in olive. Note that Tyr561 separates the canonical pore and the intracellular aqueous pathway. Glu_gate_ is located at the centre of the canonical pore where the water chain is interrupted. D. Average pore radius along canonical pores. The radius was measured using CAVER(Chovancova et al., 2012) every 1 ns during 30 ns at 0 mV. Deep and light blue colours correspond to pores 1 and 2 shown in panel B. E. Close-up showing stabilization of Glu_gate_ by interactions with residues _465_GAF_567_ and Lys212 (sticks). The CLC-2 gate is formed by Tyr561-H_2_O-Glu_gate_. Hydrogen bonds (dashed yellow lines) link Glu_gate_ to Tyr561 through one (right) or two (left) water molecules (ball and stick representation). Additionally, residues Lys568 and Glu174 on the intracellular side and Lys212 on the extracellular side are shown.

To determine the radius and shape of the pore, CLC-2^CLC-K^ and CLC-2^CLC-1^ were embedded in DMPC and POPC bilayers, hydrated with 140 mM NaCl and equilibrated using the CHARMM-GUI protocol(Allouche, 2012) at 0 mV during 100 ns and 30 ns, respectively. The initial position of Glu_gate_ in each model was different but reached the same position after equilibration (Supplementary Fig. 1). The canonical CLC-2 pore is formed by residues _164_APQAVGSGIPEM_175_, _209_PLGKEG_214_, _458_TTIPVPCGAFMP_469_, and _561_YDSIIRIK_568_ located at the N-termini of αD, αF, αN, and αR helices (Fig. 1B), respectively. Lys212 and Lys568 + Glu174 were located at the extracellular and intracellular entrances, respectively (Fig. 1E and 1F). Each monomer has a curvilinear permeation pathway filled with water and disrupted at the centre by Glu213 (Glu_gate_) (Fig. 1C). At each end of these canonical pores, there is a large water cavity. A third water cavity resembling the bifurcated pathway of hCLC-1(Park and MacKinnon, 2018) was observed in the cytosolic side (Fig. 1C, yellow star). However, this cavity is uncoupled from the canonical pore by bulky Tyr561. The radius of the pore in each monomer is shown in Fig. 1B (dark and light blue); each pore is less than 0.5 Å at the side chain of Glu_gate_ (Fig. 1D). Thereafter, the radius increases to about 2.5 Å at ± 15 Å away from Glu_gate_ and even more at both pore entrances. At 0 mV, the charged carboxyl group of Glu_gate_ is stabilized by hydrogen bonds (H-bonds) with the amino groups of _466_AF_467_ and _212_KEG_214_ (Fig. 1E). Also, Glu_gate_ forms an ionic interaction with the –OH group of Tyr561 via H-bonds mediated by one H_2_O molecule (Fig. 1F). This conformation is similar to that of Glu_gate_ in CmCLC and in rCLC-2 models built using the CmCLC structure(Feng et al., 2010; McKiernan et al., 2020). We propose that the Tyr561-H_2_O-Glu_gate_ array forms the gate that maintains CLC-2 in the closed state.

### Gating is a multistep process triggered by Cl^−^

To investigate the conformational changes that lead to CLC-2 opening we performed MD simulations using CLC-2^CLC-K^ and the three compartments/two-bilayer computational electrophysiology method (CompEl)(Kutzner et al., 2011). Each bilayer containing one channel was simultaneously exposed to the same voltage but opposite polarity during 480 ns (^+^CLC-2^CLC-K^ in bilayer 1 to +94 and +940 mV and ^−^CLC-2^CLC-K^ in bilayer 2 to −94 and −940 mV). This protocol was followed by applying a uniform electric field(Crozier et al., 2001; Roux, 2008) of −500 mV perpendicular to bilayer 2 during 200 ns.

First, we determined whether the closed Glu_gate_ position was altered by hyperpolarization or depolarization in Cl^−^-free pores (pore 2 of ^−^CLC-2^CLC-K^ and pores 1 and 2 of ^+^CLC-2^CLC-K^). Relative to 0 mV, we observed little o no changes in the position of Tyr561, Glu_gate_, and Lys212 at positive or negative voltages (Fig. 2). Regardless of voltage, Glu_gate_ was stabilized by H-bonds formed at different times with either the -OH group of Tyr561 via one H_2_O molecule (97% of the total simulation time; lifetime 63-116 ps) or with -NH group of residues _466_AF_467_ and _212_KEG_214_ (3%, lifetime 2.9-4.4 ps), and by direct interaction with Tyr561 (0.12%). When a +940 mV was applied to ^+^CLC-2^CLC-K^, the Glu_gate_ in pore 1 was pushed inwardly and displaced the water molecule that bridges Tyr561 and Glu_gate_ (Fig. 2A, cyan). This displacement increased the Tyr561-Glu_gate_ direct interaction to 40%. Thus, voltage alone cannot open the CLC-2 gate.

**Fig. 2.**
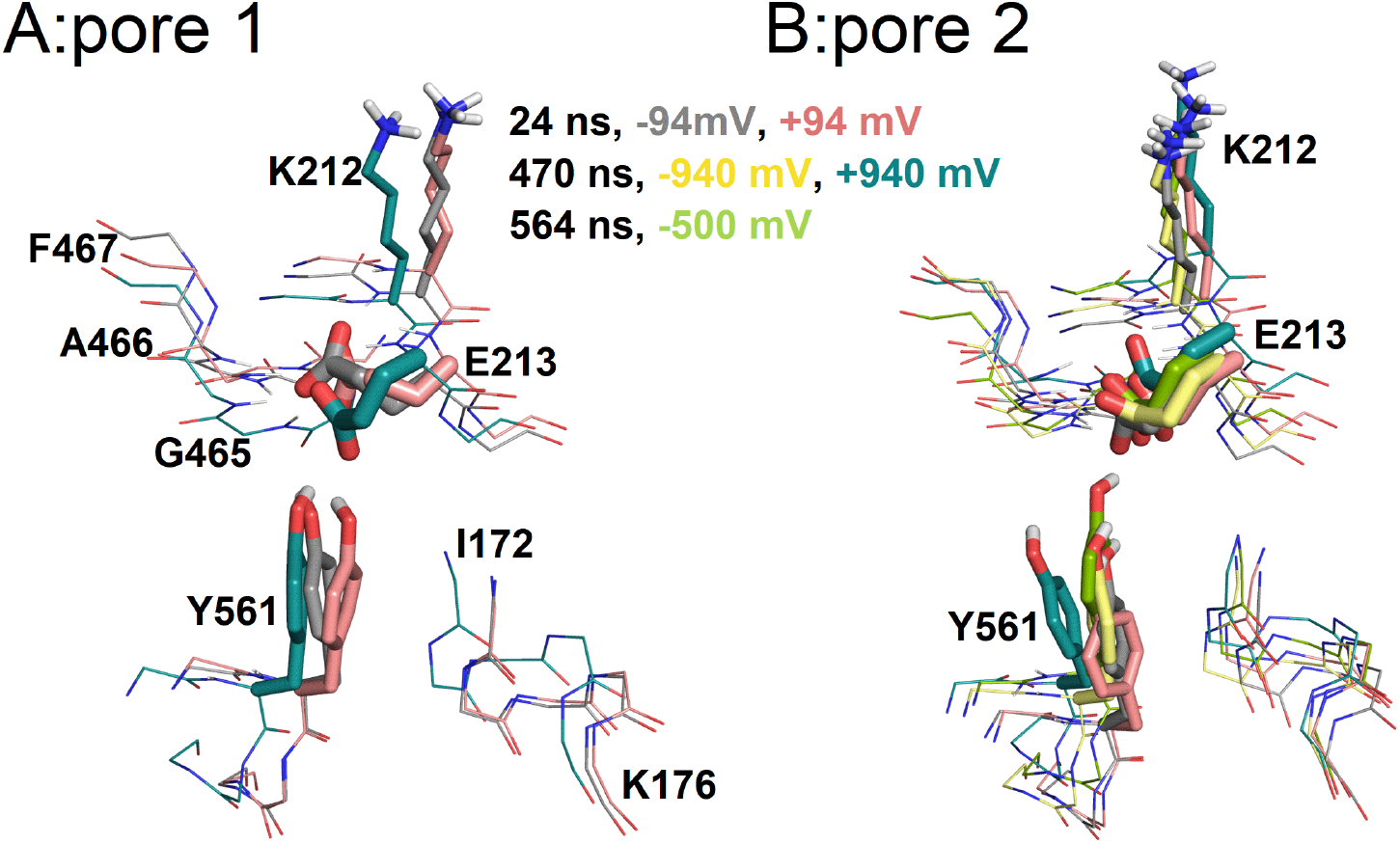
Changes in the transmembrane voltage do not trigger gating when the pores are empty. The conformation of three pores: pore 1 of ^+^CLC-2^CLC-K^ (A) and pore 2 of ^−^CLC-2^CLC-K^ and ^+^CLC-2^CLC-K^ (B) channels are depicted at different voltages (−94, −940, 500, +94, and +940 mV) and MD simulation times (24, 470, and 564 ns). The spatial position of Tyr561, Glu_gate_, and Lys212 is shown. Note that Glu_gate_ remained at the same position regardless of the transmembrane voltage.

Second, we examined the structural rearrangements that residues lining pore 1 of ^−^CLC-2^CLC-K^ underwent at negative voltage when five Cl^−^ ions occupied the pore (Cl^−^1-Cl^−^5). Based on the time (≥ 1 ns) that a Cl^−^ ion spent on sections along the pore, we defined four putative binding sites: intracellular S_i_ formed by Lys568 and Glu174, central S_c_ formed by Tyr561 and Ile172, external S_e_ formed by _465_GlyAlaPhe_467_, and exit S_o_ formed by Lys212 and Glu213. Figure 3A shows snapshots of pore 1 of ^−^CLC-2^CLC-K^ at 0, 24, 37, 470, 513, 526, 541, 564, 579, 605, 654, and 663 ns into the MD simulation (arrowheads in the site occupation roster, right side of Fig. 3B). The snapshots depict the journey of Cl^−^3 through the entire pore in five steps:

**Fig. 3.**
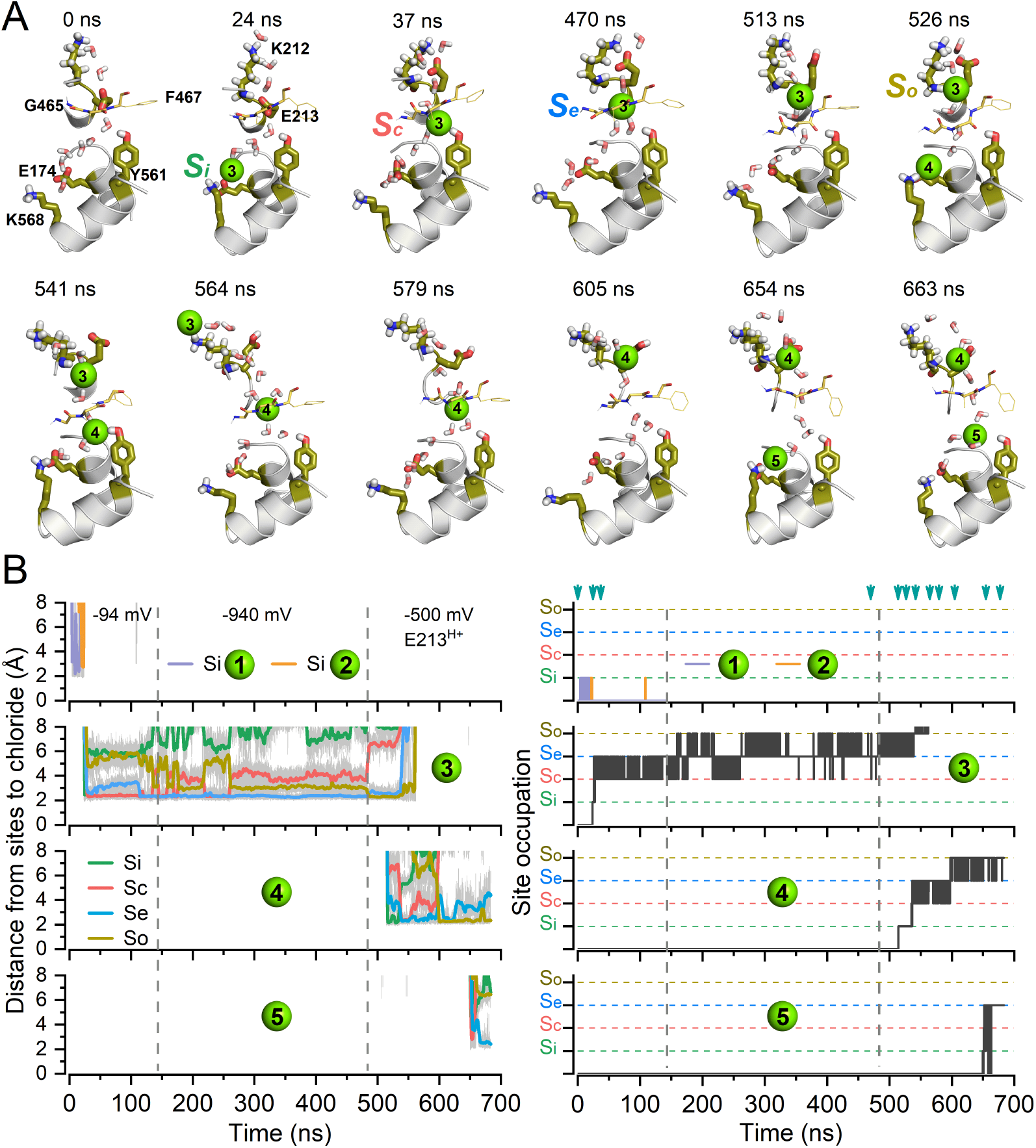
Coupling gating to permeation enables CLC-2 activation at the molecular level. A. Snapshots of structural rearrangements of key residues located in pore 1 of ^−^CLC-2^CLC-K^ channel sampled by MD simulation at the times indicated on top of each structure. Ball and stick representation of Lys568, Glu174, Tyr561, Glu213, Gly465, Ala466, Phe467, and Lys212 along with eight water molecules illustrate the canonical permeation pathway (snapshots at 0 and 24 ns). Green spheres represent Cl^−^ ions. Hyperpolarisations applied were −94, −940, and −500 mV. Snapshots are representative of pore occupation (24 ns), displacement of Glu_gate_ (37 and 470 ns), Glu_gate_ protonation (513 ns), double occupancy (526 and 541 ns) and exit of Cl^−^ from the pore (564 and 579 ns). Snapshots at 605, 654, and 663 ns illustrate changes in permeation caused by double occupancy. Residues participating in Cl^−^ transport and gating (Glu174, Lys568, Tyr561, Glu213, and Lys212) are shown in olive. The internal (Lys568-Glu174, green), centre (Tyr561, pink), external (_465_GAF_467_, blue), and extracellular (_212_KGlu_213_, olive) sites that are occupied by Cl^−^ along the pore are labelled as S_i_, S_c_, S_e_, and S_o_, respectively. B. Left: The distance separating Cl^−^ from the binding sites (S_i_, S_c_, S_e_, and S_o_) is shown as a function of the simulation time. Right: Site occupation determined by the time spent by Cl^−^ at each binding site during the simulation. Five Cl^−^ ions visited the intracellular side of the pore during the entire 680 ns MD simulation. Of these, only Cl^−^3, Cl^−^4 and Cl^−^5 occupied the pore. Cl^−^1 (lavender) and Cl^−^2 (light green) at the top, next Cl^−^3, then Cl^−^4, and Cl^−^5 at the bottom. Dashed vertical lines indicate the time when the transmembrane voltage was changed from - 94 to −940 mV and from 940 to −500 mV. −94 and −940 mV was applied using CompEl. −500 mV was applied using an electric field after Glu213 was protonated (E213^H+^). Cyan arrowheads on the top (right panel) indicate times at which the snapshots shown in A were captured.

#### Pore occupation

Figure 3A (snapshot at 0 ns and −94 mV) shows pore residues and 8 water molecules to indicate an empty permeation pathway closed by Tyr561-H_2_O-Glu_gate_ gate. The side chains of Lys568 and Glu174 fluctuated between 2-5 Å. Figure 3B shows that Cl^−^1 and Cl^−^2 occupied S_i_ for 11.5 and 3.4 ns and then spontaneously returned to the intracellular side. As Cl^−^ ions visited the pore, the side chains of Lys568 and Glu174 became closer (2 Å). Then Cl^−^3 occupied S_i_ during 3.5 ns (Figs. 3A and 3B, snapshot at 24 ns).

#### Disruption of Tyr561-H_2_O-Glu_gate_

Cl^−^3 moved to S_c_ (snapshot at 37 ns) disrupting the water bridge between Tyr561 and Glu_gate_ and binding to Tyr561. The electrostatic and steric forces (or electro-steric forces) repelled Glu_gate_ outwardly. Subsequently, Glu_gate_ interacted with the Lys212 side chain and the _213_EG_214_ backbone. Unlike in the empty pore, now Glu_gate_ stopped interacting with Gly214, folded in itself and remained attached to its backbone 59% of the time (lifetime 12.5 ps). Simultaneously, the number of H-bonds between Glu_gate_ and Lys212 increased from 1 to 3 during 98.8% of the simulated time (CompEl protocol; lifetime 274 ps). To accelerate the permeation process, we increased the voltage from −94 to −940 mV at t = 145 ns and this stabilized Cl^−^3 at S_e_ (Fig. 3B; snapshot at 470 ns). The majority of the time Cl^−^3 dwelt between S_c_ (133 ns) and S_e_ (320 ns) (Fig. 3B) and S_e_ became closer to S_o_ because _212_LysGlu_gate_ adopted an intermediate position between the closed and open configurations (Supplementary Fig. 2A).

#### Glu_gate_ protonation

We had reported that Glu_gate_ is stabilized in the open conformation by extracellular H^+^ in a voltage-independent manner(Sánchez-Rodríguez et al., 2012). Hence, we asked whether protonation was necessary to disentangle Glu_gate_ from S_e_. At t = 484 ns (Fig. 3A), the carboxyl group of Glu_gate_ was protonated using CHARMM-GUI. After protonation (snapshot at t = 513), the H-bond between Glu_gate_ and Lys212 was lost; Cl^−^3 stayed for 10.5 ns on S_e_ and then occupied S_o_ (snapshot at 526 ns).

#### Multi-ion pore occupancy

After disengaging Glu_gate_ from Lys212, Cl^−^4 entered and interacted with S_i_ for 22 ns (snapshot at 526 ns). Then Cl^−^4 jumps (snapshot at 541 ns) and fluctuates between S_c_ (for 14.4 ns) and S_e_ (57.1 ns). The hop of Cl^−^4 into S_c_ tilts Lys212 counter-clockwise by about 45° without altering Glu_gate_.

*Pore exit*. Further outward movement of Cl^−^4 near S_o_ induced straightening of Lys212 and placed Cl^−^3 onto the Lys212 side chain end for 66.7 ns (Fig. 3A, snapshot at 564 ns). The total conformational change translated the _212_KEG_214_ backbone ~6 Å away from the closed position (Supplementary Fig. 2B and Fig. 2C). The lengthy interaction of Cl^−^3 with S_o_ could be explained by an enhancement of the electrostatic potential at Lys212 position in the open state (Supplementary Fig. 3). Finally, Cl^−^3 escapes the pore (snapshot at t = 579 ns) ending its 580 ns journey while Cl^−^4 sits onto S_o_ for 76 ns (snapshot at t = 605 ns). At this time, Cl^−^5 occupies S_i_ (for 1.1 ns, snapshot at 654 ns), then S_c_ (for 4.5 ns, t = 663 ns), and S_e_ (for 27.4 ns) signalling that the pore is fully open/conductive and that Cl^−^4 is near to exit the pore.

These steps, stitched in the video, agree with previous results emphasizing that CLC-2 gating requires intracellular Cl^−^, protonation by external H^+^, and multi-ion occupancy(De Jesús-Pérez et al., 2016; Sánchez-Rodríguez et al., 2012, 2010).

### Video: 1 CLC-2 Activation mechanism

*Trajectory of Cl^−^3 along the pore at −94 mV (0-47 ns) and after protonation of Glu_gate_ at −500 mV (485-660 ns). Pore 1 of ^−^CLC-2^CLC-K^ is shown. Membrane surface profile (phosphates) is represented by grey spheres. Lys568, Glu174, Tyr561, Glu213, Gly465, Ala466, Phe467, and Lys212 are represented in olive sticks, Cl^−^ in green spheres, and helix in white. Top side = extracellular side; lower side = intracellular side.*

### Tyr561-H_2_O-Glu_gate_ array controls gating

Earlier studies with CLC-0 and CLC-1 carrying a Tyr mutation in the pore showed no change in conductance between WT and mutant channels(Accardi and Pusch, 2003; Estévez et al., 2003; Ludewig et al., 1996). Those studies led to the conclusion that Tyr561 plays no role in CLC-2 gating(McKiernan et al., 2020). However, our MD simulations show otherwise. We revealed the role of Tyr561 by mutagenesis and functional studies. The activity of Tyr561Phe, Tyr561Ala and Glu213Asp mutant channels was determined using both patch-clamp and cut-open oocyte voltage-clamp techniques. The Tyr561Ala mutant channel shows currents in both directions revealing an open channel phenotype (Fig. 4A) while WT and Tyr561Phe channels displayed currents with similar kinetics. Similarly, we reasoned that the Tyr561-H_2_O-Asp213 array in the Glu213Asp mutant loses stability and mimics the gating behaviour of Tyr561Ala. Indeed, positive voltages activate the Glu213Asp channel but negative voltages induce currents similar to those of WT; tail currents were smaller and faster.

**Fig. 4.**
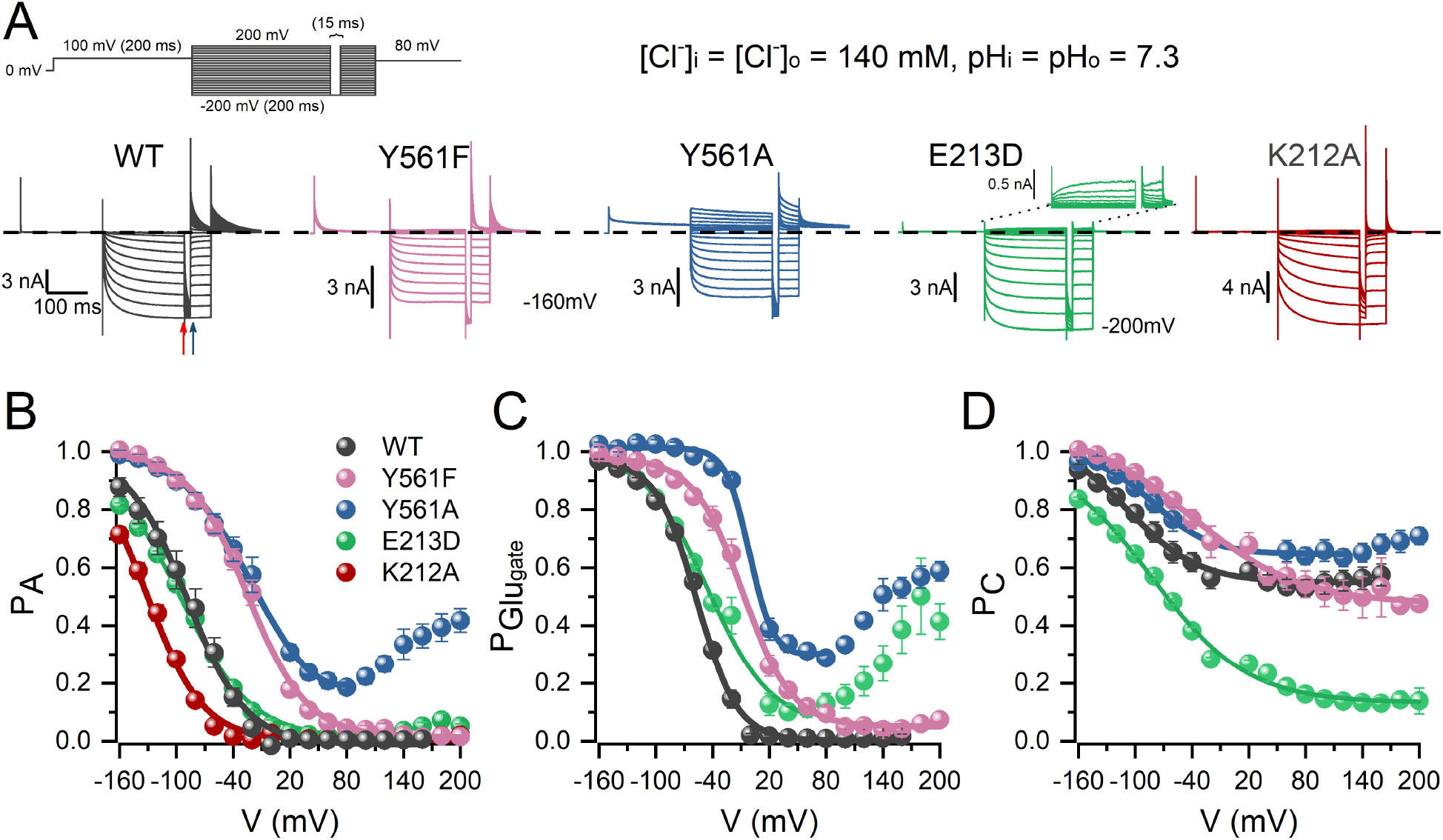
Tyr561 and Glu213 control voltage-dependent activation. A. Cl^−^ currents (colour-coded) recorded from 5 different HEK293 cells expressing WT CLC-2 (WT), Tyr561Phe (Y561F), Tyr561Ala (Y561A), Glu213Asp (E213D), and Lys212Ala (K212A). Cl^−^ currents were elicited by the voltage protocol shown in the upper panel. The protocol consisted in depolarization steps from −200 to 200 mV in 20 mV increments; we introduced an inter-pulse of 15 ms at −200 mV to reach P_Glugate_ = 1 and a repolarization voltage to +80 mV to record tail currents. Cl^−^ currents were acquired using pH_i_ = pH_o_ = 7.3 and [Cl^−^]_i_ = [Cl^−^]_o_ = 140 mM. Dash line indicates I_Cl_ = 0. B. Apparent open probability (P_A_) computed for WT CLC-2, Tyr561Phe, Tyr561Ala, Glu213Asp, and Lys212Ala. Continuous lines represent data fitted with the Boltzmann equation to determine voltage-dependent parameters V_0.5_ and z, listed in Table 1. C.,D. Voltage dependence of Glu_gate_ (P_Glugate_) and common gate (P_C_) obtained for WT CLC-2, Tyr561Phe, Tyr561Ala, Glu213Asp, and Lys212Ala. The probability of each gate was calculated as described in the methods section. Continue lines are fits to Boltzmann equation to determine the voltage-dependent parameters (V_0.5_ and z) listed in Table 1.

Values for the parameters (V_0.5_ and z) describing the voltage dependence of apparent open probability (P_A_), Glu_gate_ open probability (P_Glugate_), and common gate open probability (P_C_)(De Santiago et al., 2005) (Figs. 4B, 4C, and 4D) are summarized in Table 1. The V_0.5_ (mV)/z values for P_A_, of WT and Glu213Asp channels were quite similar: −89.3 ± 8.5/-0.84 ± 0.04 and −97.8 ± 3.3 /-0.64 ± 0.01, respectively. In contrast, the V_0.5_ values for both Tyr561Phe and Tyr561Ala mutants displayed a large rightward shift without changing z (Fig. 4B, Table 1). The changes mainly involved P_Glugate_ V_0.5_ values (Fig. 4C). For Tyr561Phe and Tyr561Ala, the P_Glugate_ V_0.5_ values were rightward shifted by ~+50 mV relative to WT P_Glugate_ (−56.6±1.0 mV). Notable, although both Glu213Asp and Tyr561Ala displayed reopening of “Glu_gate_” at positive voltages, the V_0.5_ value for Glu213Asp was similar to that of WT. Overall, the voltage dependence of P_C_ (Fig. 4D) seems to be quite similar in WT, Tyr561Phe, and Tyr561Ala. However, the voltage dependence of P_C_ in Glu213Asp indicates that the common gate starts to close at voltages that are more negative and is fully closed at positive voltages. These results indicate that Tyr561 keeps Glu_gate_ closed; a conclusion further supported by WT, Tyr561Phe, and Tyr561Ala data collected using the cut-open oocyte technique (Supplementary Fig. 4 and Table 1).

**TABLE 1.** Effect of [H^+^]_i_ and [Cl^−^]_O_ on the voltage dependent parameters of WT and mutants CLCe 2 channels. 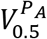 *= voltage to reach an open probability of 0.5,* 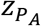 *= apparent charge of the opens closed transition,* 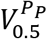 *= voltage to reach a 0.5 open probability of Glu_gate_,* 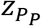 *= apparent of Glu_gate_ charge,* 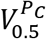 *= voltage to reach a 0.5 open probability of the common gate,* 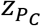 *= apparent charge the common gate. These parameter values were calculated by fitting the data to the Boltzmann equation (Equation 1). Er = reversal potentials determined by currents voltage relationship interpolations. HEK = data collected from HEK 293 cells; OOC = data collected from cut-open oocytes.*

| Channel | $V_{0.5}^{P_A}$ (mV) | $z_{P_A}$ | $V_{0.5}^{P_P}$ (mV) | $z_{P_P}$ | $V_{0.5}^{P_C}$ (mV) | $z_{P_C}$ | Er (mV) | pH<sub>i</sub> | [Cl<sup>-</sup>]<sub>o</sub> (mM) | n |
| --- | --- | --- | --- | --- | --- | --- | --- | --- | --- | --- |
| WT-HEK | -89.3 ± 8.5 | -0.84 ± 0.04 | -56.6 ± 1.0 | -1.17 ± 0.07 | -110.9 ± 6.3 | -0.75 ± 0.03 |  | 7.3 | 140 | 5 |
| WT-OOC | -96.9 ± 7.5 | -0.87 ± 0.04 | -72.3 ± 1.4 | -1.24 ± 0.10 | -115.1 ± 5.5 | -1.17 ± 0.13 |  | 7.3 | 140 | 7 |
| WT | -79.1 ± 5.8 | -0.79 ± 0.07 | -39.5 ± 2.2 | -1.19 ± 0.07 | -82.9 ± 6.1 | -0.54 ± 0.04 | 57.4 ± 0.6 | 7.3 | 10 | 5 |
| Y561F-HEK | -34.8 ± 9.4 | -0.8 ± 0.0 | -13.8 ± 6.4 | -1.1 ± 0.1 | -51.2 ± 6.4 | -0.4 ± 0.0 |  | 7.3 | 140 | 9 |
| Y561F-OOC | -29.7 ± 7.6 | -0.57 ± 0.05 | -5.66 ± 9.9 | -0.71 ± 0.05 | -37.3 ± 17.9 | -0.62 ± 0.1 |  | 7.3 | 140 | 7 |
| Y561F | -68.9 ± 5.4 | -0.67 ± 0.02 | -6.3 ± 4.4 | -0.73 ± 0.04 | -71.9 ± 4.5 | -0.62 ± 0.07 | 52.5 ± 1.6 | 7.3 | 10 | 6 |
| Y561F | -40.0 ± 4.5 | -0.77 ± 0.04 | -21.6 ± 2.2 | -0.95 ± 0.08 | -36.1 ± 7.9 | -0.53 ± 0.06 |  | 5.5 | 140 | 8 |
| Y561F | -60.3 ± 17.8 | -0.73 ± 0.08 | -17.9 ± 8.6 | -1.09 ± 0.17 | -78.7 ± 23.2 | -0.63 ± 0.06 |  | 4.3 | 140 | 5 |
| Y561A-HEK | -30.4 ± 5.9 | -0.84 ± 0.09 | -4.1 ± 2.5 | -1.48 ± 0.12 | -65.6 ± 10.6 | -0.81 ± 0.06 |  | 7.3 | 140 | 7 |
| Y561A-OOC | -37.4 ± 6.1 | -0.85 ± 0.04 | -41.8 ± 15.2 | -1.18 ± 0.08 | -64.0 ± 1.88 | -1.05 ± 0.01 |  | 7.3 | 140 | 3 |
| Y561A | -58.6 ± 8.6 | -0.63 ± 0.01 | 25.8 ± 4.7 | -1.08 ± 0.06 | -74.5 ± 7.3 | -0.64 ± 0.07 | 57.5 ± 0.8 | 7.3 | 10 | 6 |
| Y561A | -31.7 ± 6.5 | -0.82 ± 0.07 | -13.4 ± 2.6 | -1.04 ± 0.08 | -77.2 ± 15.9 | -0.98 ± 0.16 |  | 5.5 | 140 | 9 |
| Y561A | -38.6 ± 8.0 | -0.89 ± 0.15 | -12.7 ± 5.2 | -1.26 ± 0.09 | -66.7 ± 12.1 | -0.77 ± 0.07 |  | 4.3 | 140 | 10 |
| E213D | -97.8 ± 3.3 | -0.64 ± 0.01 | -44.6 ± 3.6 | -0.82 ± 0.05 | -86.0 ± 6.1 | -0.51 ± 0.01 |  | 7.3 | 140 | 8 |
| K212 | -130.7 ± 3.6 | -1.16 ± 0.03 | -- | -- | -- | -- | -- | 7.3 | 140 | 6 |

MD simulations suggest that Lys212 is relevant for Cl^−^ exit. To confirm the role of Lys212, we first calculated the electrostatic potential in the closed and open pores of WT and ^−^CLC-2^CLC-K^ Lys212Ala mutant (produced *in silico*). The electrostatic potential decreased from −26 Kcal/mol to −12 Kcal/mol (closed) and from −52 Kcal/mol to −40 Kcal/mol (open) after mutating Lys212 (Supplementary Fig. 3). Then, we determined the voltage dependence of P_A_ from whole cell recordings of the Lys212Ala mutant channel (Fig. 4A). Mutant channels activated with kinetics similar to that of WT; at −200 mV, the time constants of the fast kinetic component were 3.2 ± 0.7 ms and 3.8 ± 0.5 ms, for the slow component were 41.0 ± 3.9 ms and 40.1 ± 7.9 ms for mutant and WT, respectively. Unlike WT, the mutant channels had very fast closing; at +80 mV, the time constants were 4.7 ± 0.24 ms for mutant and 59.4 ± 4.8 ms for WT. This result indicates that Lys212 helps to keep the pore in the open conformation. In agreement with this idea, the voltage dependence of P_A_ in the Lys212Ala mutant channel (Fig 4B; Table 1) was shifted to negative voltages (V_0.5_/z = −130.7±3.6 mV/1.16±0.03).

WT CLC-2 has low sensitivity to extracellular [Cl^−^] ([Cl^−^]_o_)(Sánchez-Rodríguez et al., 2010). However, Glu_gate_ could become sensitive to [Cl^−^]_o_ when is dislodged, as in the Tyr561Ala mutant (Fig. 4). The Cl^−^ influx in Tyr561 mutants would knock off Glu_gate_ inwardly just as the Cl^−^ efflux knocks off the WT Glu_gate_ outwardly. Under whole cell condition, the [Cl^−^]_o_ was changed from 140 to 10 mM, this manoeuvre decreased Cl^−^ influx but had little effect on the kinetics of WT, Ty561Phe, and Tyr561Ala currents (Fig. 5A, 5B and 5C). The voltage dependence of P_A_ and P_Glugate_ in the WT channel was little or not affected by varying the [Cl^−^]_o_ (Fig. 5A, Table 1).

**Fig. 5.**
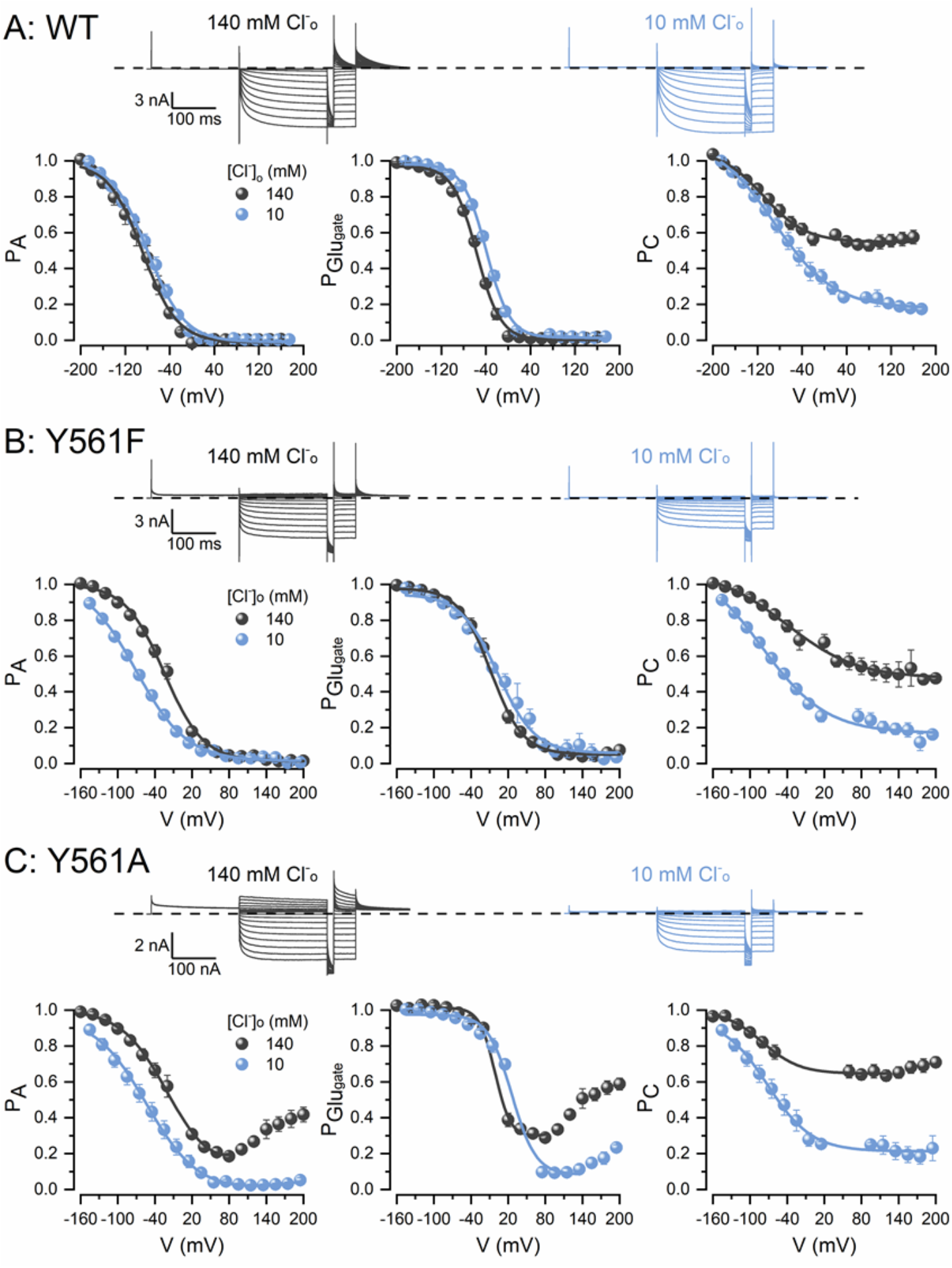
TMEM16A channels that have Tyr561 mutated to Phe or Ala are sensitive to extracellular chloride. A,B,C. Top panels: Cl^−^ currents recorded from the same HEK293 cell expressing WT CLC-2 (A), Tyr561Phe (B), or Tyr561Ala (C) exposed first to 140 (grey) and then 10 (blue) mM [Cl^−^]_o_ at pH_o,_ = 7.3. Channels were activated using the voltage protocol shown in Fig. 4A. A,B,C. Bottom panels: Voltage dependence of P_A_ (left), P_Glugate_ (middle), and P_C_ (right) for WT CLC-2 (A), Tyr561Phe (B) and Tyr561Ala (C) determined first with 140 (grey) and then with 10 (blue) mM extracellular Cl^−^. [Cl^−^]_i_ = 140 mM and pH_i_ = pH_o_ = 7.3. Lines are fits of the data with Boltzmann equation. Voltage-dependent parameters V_0.5_ and z are listed in Table 1.

However, in Tyr561Phe channels (Fig. 5B), we observed a −34 mV shift on V_0.5_ of P_A_ with little effect on P_Glugate_ (Table 1). A severe alteration of the voltage dependence of both P_A_ and P_Glugate_ was observed in the Tyr561Ala mutant at positive voltages (Fig. 5C). In this mutant, the voltage dependence of P_A_ was shifted by about −29 mV and re-opening at positive voltages was abolished by lowering the [Cl^−^]_o_ from 140 to 10 mM. Similarly, P_Glugate_ was strongly diminished at positive voltages. That is, decreasing the Cl^−^ influx decreased the open probability of Glu_gate_ at positive potentials in the Tyr561Ala mutant channel. In contrast, the effect of low [Cl^−^]_o_ on V_0.5_ of P_C_ was nearly the same in WT, Tyr561Phe and Tyr561Ala channels (V_0.5_ were between −71 to −82 mV, Table 1). The common gate tended to close at low external Cl^−^ conditions.

To determine the conformational state of Glu_gate_ when Try561 is mutated, we analysed the pore structures of Tyr561Phe and Tyr561Ala mutants made *in-silico*. In contrast to the WT CLC-2 pores, the pore structures of Tyr561Phe and Tyr561Ala show a disordered Tyr561-H_2_O-Glu_gate_ array in both pores (Supplementary Fig. 5). Although Glu_gate_ still interacted through H-bonds with the -NH groups of _466_AF_467_ and _212_KEG_214_, the Tyr561Phe-H_2_O-Glu_gate_ bridge was formed only 16% of the time whereas the bridge Tyr561Ala-H_2_O-Glu_gate_ was not formed at all. Hence, the Tyr561-H_2_O-Glu_gate_ array is critical for maintaining the pore in a closed state.

### The anomalous mole fraction behaviour of CLC-2 activation reveals the multi-ion occupancy of the pore

MD simulations show that Cl^−^ can disrupt the Tyr561-H_2_O-Glu_gate_ array and that Cl^−^ exit is facilitated when the pore is occupied by two Cl^−^ ions. These observations are in agreement with the idea that multi-ion occupancy(Hille, 2001) facilitates permeation by a knock-on mechanism(Khalili-Araghi et al., 2006; Kratochvil et al., 2016). To reveal the multi-ion occupancy of wild type CLC-2 channels we determine the anomalous mole fraction (AMF) behaviour using different acetate/Cl^−^ mole-fractions. As the acetate mole-fraction increased, we recorded less inward Cl^−^ currents and the depolarization-induced tail currents that saturated at less positive voltages (Fig. 6A). Increasing the acetate mole fraction shifted the voltage-dependent activation curves first to the right and then to the left on the voltage axis (Fig. 6B). The corresponding V_0.5_ values, when plotted against the acetate/Cl^−^ mole fractions, had a concave shape and peaked at 0.57 (Fig. 5D, black spheres). These data support the idea that multi-anion occupancy of the pore controls CLC-2 gating. If acetate and Cl^−^ occupy the pore then both anions contribute to the AMF dependence. To test this idea, we dialyzed the cells with acetate mole fractions that had the pH adjusted to 4.2 to protonate 79% of the acetate. Under this condition, we did not find evidence of AMF dependence in voltage activation (Fig. 6C). V_0.5_ had a linear relationship with acetate/Cl^−^ mole-fractions (Fig. 6D, open circles), a behaviour similar to that observed with pure Cl^−^ solutions adjusted to pH 7.3 (Fig. 6D, green squares)(Sánchez-Rodríguez et al., 2010). This result also indicates that intracellular Cl^−^ but not intracellular H^+^ control voltage activation.

**Fig. 6.**
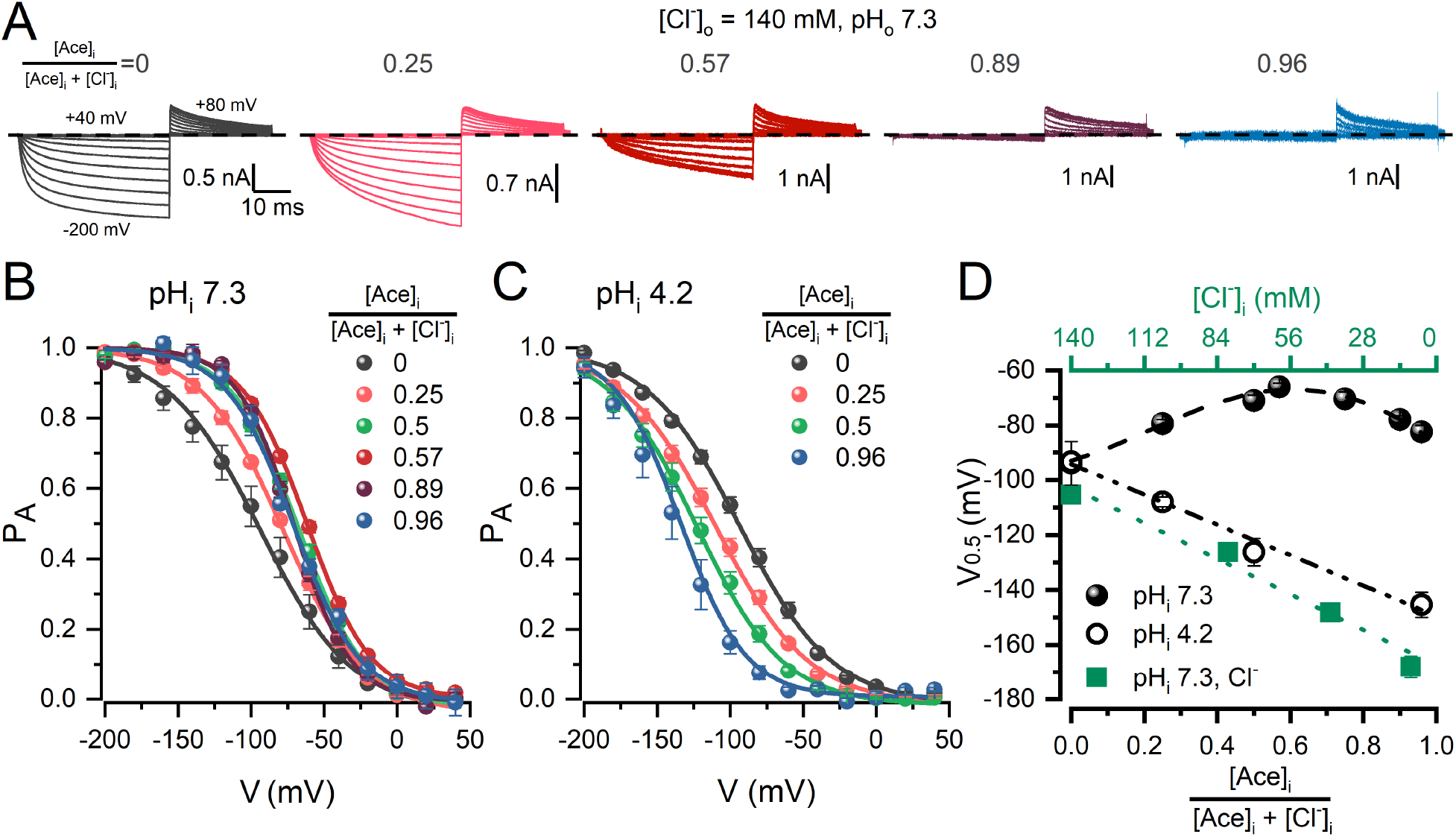
Multi-ion pore occupancy regulates voltage-dependent activation. A. Whole cell currents recorded from 5 different HEK cells expressing WT CLC-2 dialyzed with 0, 0.25, 0.57, 0.89, and 0.96 acetate mole fractions ([Ace]_i_/([Ace]_i_+[Cl^−^]_i_)). Currents were elicited by depolarization steps from −200 to +40 mV in increments of 20 mV. Tail currents were recorded at +80 mV. [Cl^−^]_o_ = 140 mM, pH_e_ = pH_i_ = 7.3. Dash line indicates the 0 current level. B,C. Voltage-dependent activation determined at pH_i_ of 7.3 (B) and 4.2 (C) using different acetate mole fractions. P_A_ was calculated using the tail current magnitude. The resulting curves were normalized to their respective tail current maximum obtained after fitting with the Boltzmann equation. Continuous lines are fits from which the voltage-dependent parameter V_0.5_ and z were computed. D. V_0.5_ plotted against acetate mole fractions (circles) and [Cl^−^]_i_ (squares). Filled symbols correspond to data obtained at pH_i_ 7.3 whereas open symbols were collected at pH_i_ 4.2. V_0.5_ values were obtained from Boltzmann fits to data shown in panels B and C. Similar data collected using intracellular solutions that contained unmixed Cl^−^ at pH_i_ = 7.3 was used to calculate V_0.5_ (green squares).

### Tyr561-H_2_O-Glu_gate_ array inhibits Glu_gate_ protonation from the intracellular side

A remarkable difference between CLC-0/CLC-1 and CLC-2 is their contrasting sensitivity to intracellular H^+^. CLC-0 and CLC-1 require Glu_gate_ protonation for activation whereas CLC-2 does not(Rychkov et al., 1996; Sánchez-Rodríguez et al., 2012; Traverso et al., 2006b). The recent structure of hCLC-1 shows a bifurcation of the permeation pathway at the cytosolic side(Park and MacKinnon, 2018). For CLC exchangers this alternative pathway enables protonation of the Glu_gate_ from the cytosolic side(Accardi et al., 2005). Our CLC-2 homology structures show the presence of an uncoupled aqueous cavity at the same position as in CLC exchangers and hCLC-1 channels (Fig. 1C). Also, the homology model shows that Tyr561, Ile172, Phe269 and Thr520, all hydrophobic residues, form a shield that faces the aqueous lacuna. We propose that these residues form a hydrophobic gate that prevents pore bifurcation. The structures of Tyr561Phe and Tyr561Ala shown in Figs. 7A and 7E, respectively, support this idea. The pore in Tyr561Phe channels is occasionally bifurcated (Fig. 7A) but it becomes permanently bifurcated only in the Tyr561Ala mutant (Fig. 7E). The bifurcation creates a forked water chain that exposes Glu_gate_ to both the alternative and the canonical pathways. The hydrophobic gates of Tyr561Phe and Tyr561Ala channels are shown in Figs. 7B and 7F, respectively, with yellow dashed lines indicating the distance between paired residues. Fig. 7D shows distances between Phe269-Tyr561 and Thr520-Ile172 in the two pores of WT CLC-2 and between Phe269-Tyr561Phe and Thr520-Ile172 in the two pores of Tyr561Phe mutant channel. The distances remained around 3 Å and 6 Å, respectively, in both pairs, except for pore 2 in Phe269-Tyr561Phe which was about 6 Å. In contrast, the equivalent distances in both pores of the Tyr561Ala mutant increased to around 5-11 Å (Fig. 7H). The consequence of these distortions in the hydrophobic gate was a widening of the alternative pore radius with minimal effects on the canonical pore radius (Fig. 7C and 7G). The radius of the alternative pore in Tyr561Ala mutant became nearly identical to the radius of the intracellular portion of the WT canonical pore (Fig. 1D). The alternative pore of Tyr561Ala is wider than that of Tyr561Phe, which allowed the formation of the forked water chain.

**Fig. 7.**
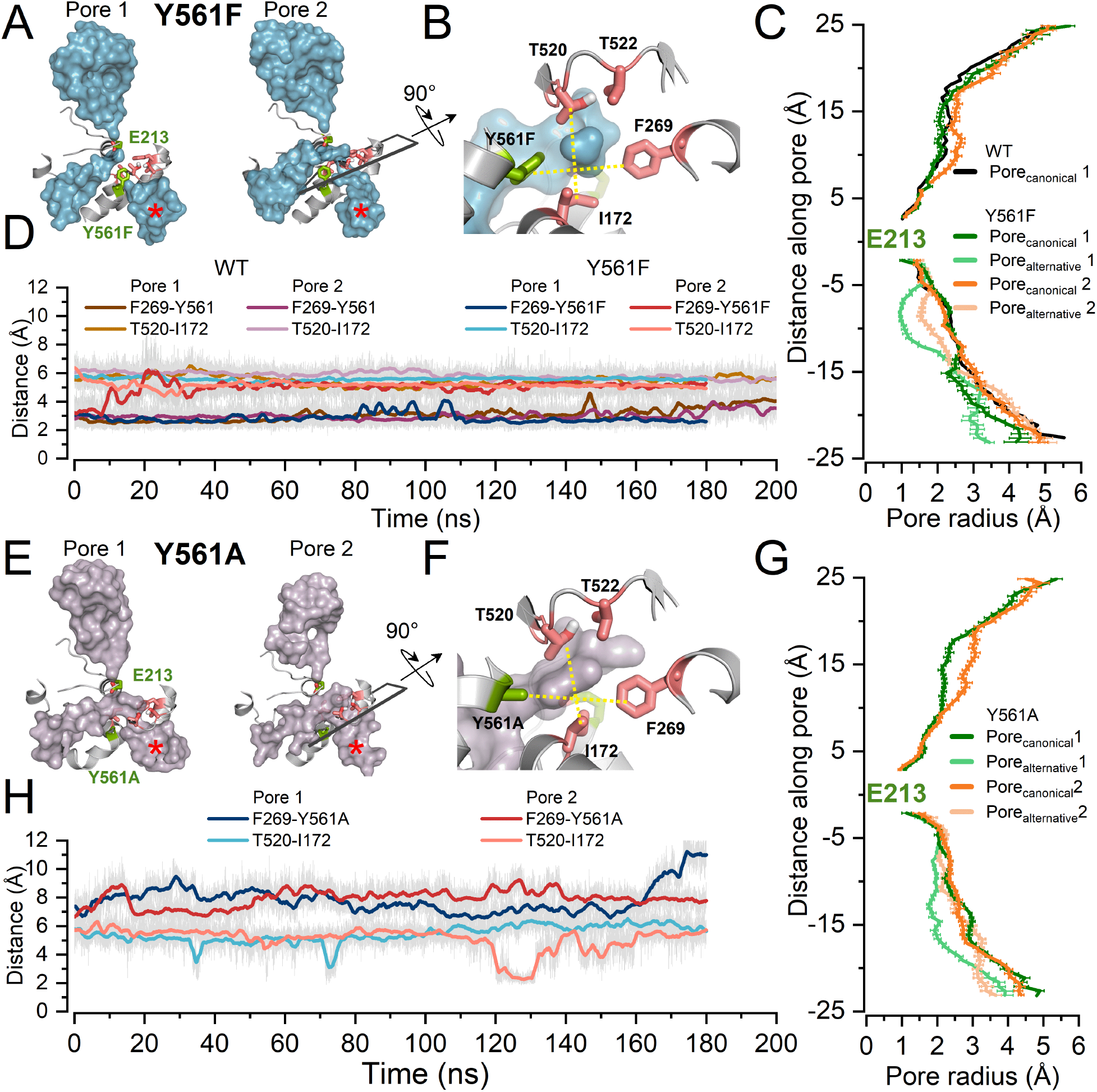
The canonical pores of CLC-2 are uncoupled from an alternative pathway leading to the cytosol via a hydrophobic gate formed by Tyr561, Ile172, Phe269, and Thr520. A,E. Canonical pores and the alternative pathways (indicated by red asterisks) are represented by the surface of water molecules in Tyr561Phe (A) and Tyr561Ala (E) mutants after 180 ns MD simulation at −500 mV. Residues Tyr561Phe, Tyr561Ala, and Glu_gate_ are coloured in green. The tilted rectangles in pore 2 of both Tyr561Phe and Tyr561Ala mutants indicate the location plane for residues Ile172, Phe269, and Thr520 (pink) that comprise part of the hydrophobic gate. B,F. Residues Ile172, Phe269, Thr520, Thr522, and Tyr561Phe (B) or Tyr561Ala (F) create a hydrophobic gate. Magnification of the planes delimited by black rectangles in A and E are rotated 90° to show the hydrophobic gate. The distances between residues connected by yellow dashed lines were measured to determine the size of the hydrophobic gate in Tyr561Phe (B) and Tyr561Ala (F) mutant channels. D,H. Distances between residues of the hydrophobic gates in WT, Tyr561Phe, and Tyr561Ala channels as a function of the simulation time. Distances were measured diagonally between two pairs of residues: Phe269-Tyr561 and Thr520-Ile172 in WT, Phe269-Tyr561Phe and Thr520-Ile172 in the Tyr561Phe mutant (D) and between Phe269-Tyr561Ala and Thr520-Ile172 in the Tyr561Ala mutant (H). Measurements made at −500mV. C,G. Average pore radius along canonical and alternative pores in WT and Tyr561Phe (C) and Tyr561Ala (G) mutants. The radius of both pores was measured and plotted for each channel. Canonical pores: 1 = black and dark green, 2 = dark orange; alternative pores: 1 = light green, 2 = light orange. The radius was measured at 0 mV during 30 ns every 1 ns. Note that the alternative pore is located after Glu_gate_, between 0 and −25 Å.

We speculate that protonation of CLC-2 Glu_gate_ by intracellular H^+^ is prevented(Sánchez-Rodríguez et al., 2012) by the hydrophobic gate that precludes pore bifurcation in WT CLC-2. To verify this we studied voltage-dependent activation at different intracellular pH (7.3, 5.5, and 4.3) in Tyr561Phe and Tyr561Ala mutants that have partially and fully bifurcated pores. Fig. 8 shows that both mutants were readily accessible to intracellular H^+^. Tyr561Ala channels show larger currents at positive potentials than Tyr561Phe channels upon intracellular acidification (Figs. 8A and 8B). Relative to pH_i_ 5.5 the voltage dependence (V_0.5_) of Tyr561Phe P_A_ (Fig 8A) at pH_i_ 4.3 was reduced by 20 mV, whereas V_0.5_ for Tyr561Ala (Fig. 8B) was reduced by 7 mV (Table 1). Notably, in Tyr561Ala the effects were readily measured at pH_i_ 5.5 and P_A_ increased from 0.2 to nearly 0.6 after decreasing pH_i_ from 7.3 to 4.3 (Fig. 8B). Similar effects were observed in the voltage dependence of Glu_gate_ in mutants Tyr561Phe and Tyr561Ala (Figs. 8A and 8B). At pH_i_ 4.3, Glu_gate_ of Tyr561Phe showed a reopening pattern at positive voltages whereas Glu_gate_ of Tyr561Ala showed stronger activation at pH_i_ 5.5 and 4.3. At pH_i_ 4.3, P_Glugate_ reached a minimum value of about 0.7 indicating that Glu_gate_ remains nearly always open at this pH_i_. Despite the reopening at positive voltages induced by intracellular H^+^, the V_0.5_ of P_Glugate_ was slightly altered in both mutants at negative potentials (Table 1). Intracellular acidification also changed the magnitude and voltage dependence of P_C_. The V_0.5_ value for Tyr561Phe was shifted by −42 mV at pH_i_ 4.3 relative to pH_i_ 5.5, and by −10 mV for Tyr561Ala (Table 1).

**Fig. 8.**
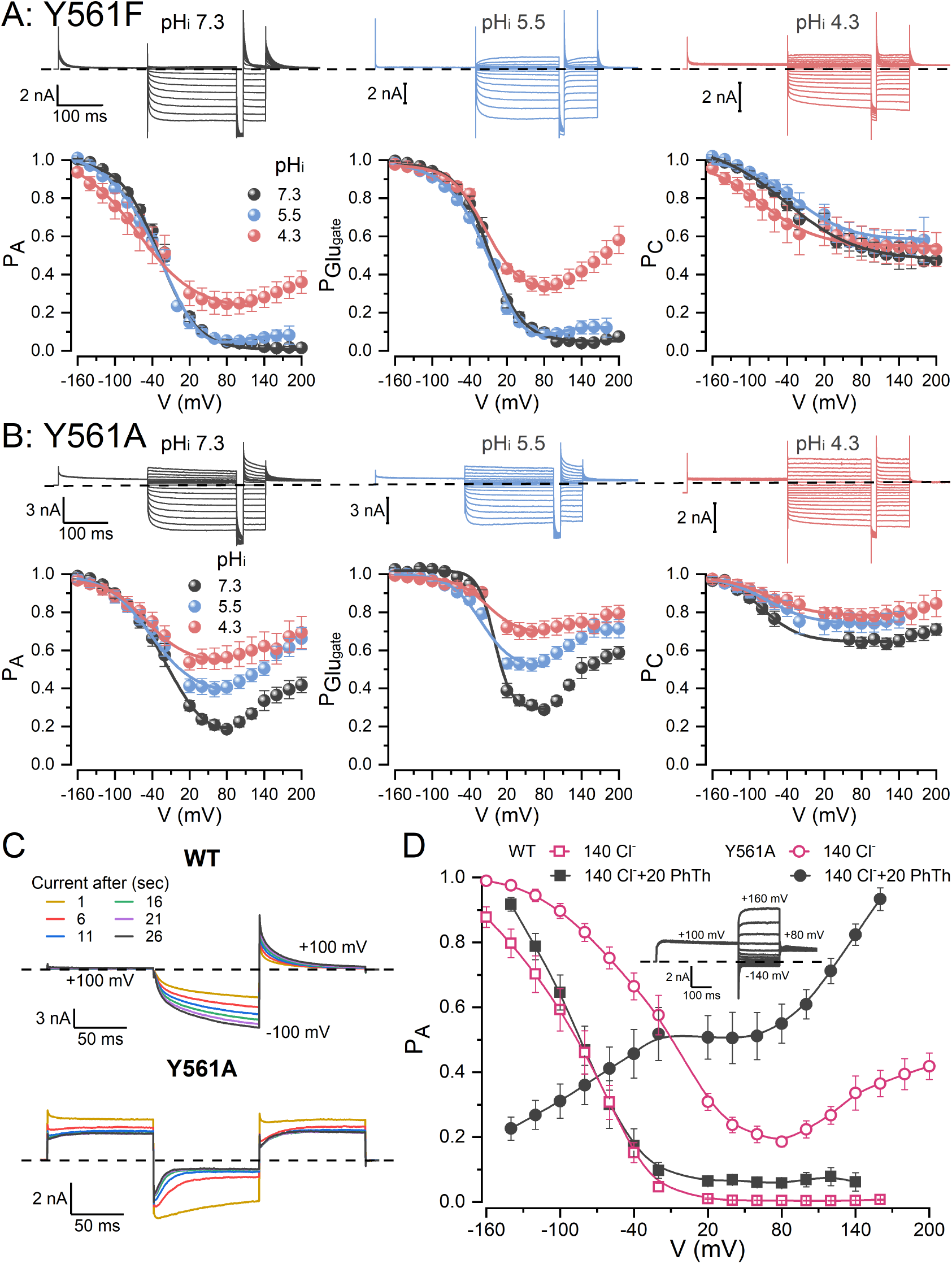
Unrestrained Glu_gate_ renders CLC-2 sensitive to intracellular protons and phthalate. A,B. Top panels. Cl^−^ currents from 3 different HEK293 cells expressing the Tyr561Phe (A) or Tyr561Ala (B) mutants were recorded in the presence of 140 mM intracellular Cl^−^ and intracellular pH 7.3 (grey), 5.5 (blue), 4.3 (red). The voltage protocol showed in Fig. 4A was utilized in these experiments. Extracellular pH and [Cl^−^]_o_ were 7.3 and 140 mM, respectively. A,B. Bottom panels. Voltage dependence of P_A_, P_Glugate_ and P_C_ for Tyr561Phe (A) and Tyr561Ala (B) are plotted using data collected at pH_i_ 7.3 (grey), 5.5 (blue) and 4.3 (red). Lines are fits of the data with the Boltzmann equation used to calculate V_0.5_ and z parameters listed in Table 1. C. 20 mM phthalate increases the whole cell current in WT CLC-2 (top panel) but inhibits the current in the Tyr561Ala (bottom panel) mutant channel. Whole cell currents were measured every 5 s after breaking the seal to get a pseudo-basal current. D The inward rectification displayed by the WT and Tyr561Ala channels is inverted by phthalate in the Tyr561Ala channel. Comparison of the voltage-dependent activation curves determined for WT (squares) and Tyr561Ala (circles) in the absence (open) and presence (filled) of 20 mM phthalate. Phthalate was present in the intracellular solution. The inset shows whole cell recording from a cell expressing Tyr561Ala that was dialyzed with 20 mM phthalate.

Further evidence of a functional alternative pore was obtained by evaluating the effect of 20 mM phthalic acid at pH 7.3 applied to the intracellular side of WT and Tyr561Ala channels. Figure 8C shows that WT CLC-2 was not blocked by phthalic acid but Tyr561Ala mutant channels were. WT CLC-2 showed typical inwardly rectifying kinetics and the current increased during 26 s of recording. In contrast, the current through Tyr561Ala mutant was blocked as soon as the membrane was hyperpolarized and then unblocked at depolarized potential (Fig. 8C, lower panel). The voltage-dependent blockade by phthalate inverted the current rectification observed in Tyr561Ala (Insert in Fig. 8D). The current saturated at negative voltages and increased at a positive potential. Fig. 8D summarizes the voltage dependence of WT (squares) and Tyr561Ala (circles) channels in the absence (empty symbols) and presence (filled symbols) of 20 mM phthalic acid in the cytosolic side. Taken together, the structural data and the effects of intracellular H^+^ and phthalate on WT and Tyr561Ala channels, indicate that the alternative pathway is not connected to the canonical pore in WT CLC-2 but it is connected in the Ty561Ala channel. These results explain the lack of protonation by intracellular H^+^ in WT CLC-2 channels.

## Discussion

### The canonical pore stays closed by Tyr561-H_2_O-Glu_gate_ and is opened by electro-steric repulsion

The general features of the CLC proteins (TMD containing two pores, C-terminus with CB domains, and a curvilinear pore with a narrow constriction at Glu_gate_ location) are conserved in the equilibrated CLC-2 homology structure. However, there are important differences in specific regions associated with key functions. In CLC-2, Glu_gate_ is slightly bent towards the intracellular side making contact with Tyr561 via one H_2_O molecule, a conformation apt for electro-steric repulsion with anions occupying the pore. This conformation of Glu_gate_ is analogous to that in the EcCLC and CmCLC Cl^−^/H^+^ exchangers but is different from the CLC-1 channel where Glu_gate_ is oriented off the pore lumen. At depolarized voltages, the pore of CLC-2 is a single empty pathway closed by Tyr561-H_2_O-Glu_gate_ whereas the pore of CLC-1 is an open bifurcated pathway occupied by two Cl^−^ ions(Park and MacKinnon, 2018). At hyperpolarized voltages, CLC-2 becomes open and occupied by two Cl^−^ ions. The closed/empty and open/occupied configurations at positive and negative voltages, respectively, correspond with CLC-2 function. Importantly, the canonical pores in CLC exchangers and CLC-1 channel are bifurcated but not in CLC-2 channel. In CLC Cl^−^/H^+^ exchangers, the alternative cavity includes Glu203 that is critical for Glu_gate_ protonation(Accardi et al., 2005), in CLC channels this residue is replaced by valine. In contrast, the pore of CLC-2 is isolated from the alternative cavity by hydrophobic residues that make a square-shaped hydrophobic gate. When we disrupted the hydrophobic gate, the pore became bifurcated and sensitive to intracellular H^+^. We propose that a non-bifurcated pore or the absence of Glu residue or both explains why CLC-2 is insensitive to intracellular H^+^. Tyr561 successfully shields CLC-2 from intracellular H^+^.

Our data revealed that a two-leaf gate formed by the ionic interaction of Glu_gate_ with the -OH group of Tyr561 mediated by one polarized water molecule, works to close the canonical pore of CLC-2. Tyr561 is stiff and remains motionless during the entire gating process. Instead, Glu_gate_ is bound to Tyr561 in empty pores held at positive voltages but is released by Cl^−^ during activation. Free Glu_gate_ moves outwardly or inwardly depending on the electro-steric repulsive force generated by the Cl^−^ flux. Accordingly, mutating Tyr561 or Glu_gate_ leads to partially open (Tyr561Phe, Glu213Asp) or fully open (Tyr561Ala, Glu213A/V) phyenotypes(De Santiago et al., 2005; Niemeyer et al., 2003). We conclude that CLC-2 Glu_gate_ operates as a one-way check valve similar to what has been proposed for K2P potassium channels(Schewe et al., 2016) and that Tyr561 prevents Glu_gate_ from opening in the inward direction.

### Activation is possible by coupling gating to permeation

Cl^−^ permeation was coupled to gating and lasted 580 ns (see video). During this time lapse, activation took place in five steps: the occupation of the pore by intracellular Cl^−^, repulsion of Glu_gate_ by Cl^−^, protonation of Glu_gate_, double occupancy of the pore, and Cl^−^ exit (Fig. 9). During activation, major structural rearrangements occurred at the intracellular side, the gate and the extracellular side. At the intracellular side, loading and emptying S_i_ trigger a rapid interaction (8.3 ns) between the side chains of Lys568 and Glu174, similar to the opening/closing of a crab claw. At the extracellular side, a protonated Glu_gate_ stays far from Lys212, which allows the straightening and counter-clockwise bent of Lys212. This remarkably structural rearrangement allows Cl^−^ to interact with the Lys212 side chain during the exit process.

**Fig. 9.**
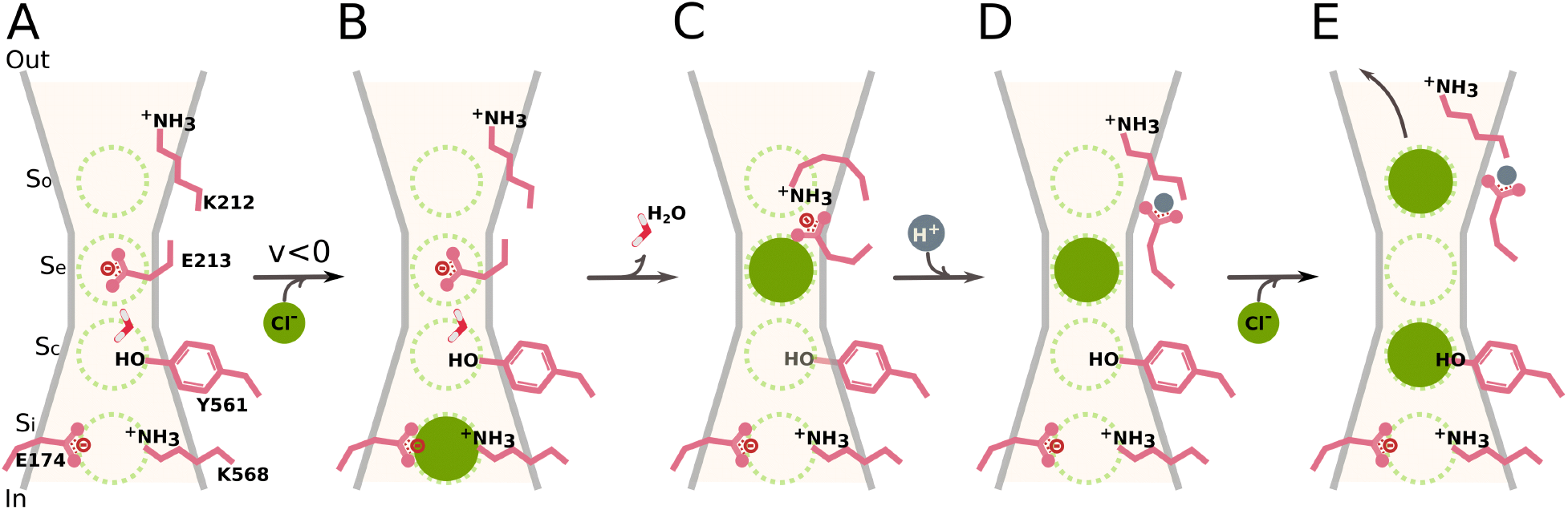
Schematic representation of the electro-steric activation mechanism for CLC-2. The Scheme depicts one pore, four anion-binding sites (S_i_, S_C_, S_e_, and S_O_), and the side chains of critical residues (Glu174, Lys568, Tyr561, Glu213, and Lys212) lining the pore. Regardless of voltage, the empty pore remains closed by the gate Tyr561-H_2_O-Glu_gate_ (A). Upon a hyperpolarization (V < 0), one intracellular Cl^−^ occupies S_i_ marking the beginning of the activation process (B). The gate is split by electro-steric repulsion when Cl^−^ moves forward into the permeation pathway and sits on S_C_ and S_e_ thus coupling permeation to gating (C). Then, Glu_gate_ adopts an outward-facing conformation and interacts with Lys212, this conformation is suggestive of a pre-open state (D). The Glu_gate_-Lys212 salt-bridge is broken by protonation allowing an outward straightening of the Lys212 side chain. Finally, Cl^−^ exits the pore when another intracellular Cl^−^ occupies S_i_ and S_C_ (E). A doubly occupied pore indicates that the pore is fully open and conductive.

Despite that, the loaded Lys212 side chain was near to extracellular media, the Cl^−^ does not readily escape because it sits in a deep well. For Cl^−^ to leave Lys212 another Cl^−^ ion needs to occupy S_i_. The doubly occupied pore condition (for 100 ns) is critical for initiating and maintaining anion conduction (Fig. 9, video). A recent computational analysis aimed at predicting intrinsic or extrinsic elements of a protein that could respond to changes in the local electric field indicated that either Glu_gate_ or a Cl^−^ ion trapped in the pore could participate in voltage sensing in CLC-1(Kasimova et al., 2018). According to our simulations, Glu_gate_ moved outwardly ~7 Å from its original closed position but remained interacting with Lys212 until it was protonated. A protonated Glu_gate_ do not contribute to voltage sensing. Instead, is the negatively charged permeant Cl^−^ ion that goes across the entire electrical field that provides the gating charge in CLC-2.

The gating mechanism proposed here differs substantially from the mechanism recently suggested by McKiernan et al,(McKiernan et al., 2020). These authors combined Markov state modelling with MD simulations and a homology structure of rat CLC-2 built using the CmCLC Cl^−^/H^+^ exchanger structure 3ORG as a template(Feng et al., 2010). They propose that entry of intracellular Cl^−^ requires rotation of the Ser168-Gly169-Ile170 backbone and opening of Glu_gate_ follows this in a Cl^−^-dependent manner. However, our results show that Lys568 and Glu174 are necessary for pore occupation by intracellular Cl^−^. Another discrepancy is the role of Tyr561. McKiernan’s work suggests that Tyr561 (Tyr559 in their homology structure) is irrelevant for CLC-2 gating, an idea supported by the lack of effect on single-channel conductance after mutating Tyr in CLC-0 and CLC-1 channels(Accardi and Pusch, 2003; Estévez et al., 2003; Ludewig et al., 1996). We observed that Glu_gate_ and Tyr561 remained in the same position regardless of voltage in empty pores. But, when a pore became occupied at a negative potential, Cl^−^ rapidly reconfigured the Tyr561-H_2_O-Glu_gate_ array without disturbing Tyr561. These observations would suggest that Tyr561 does not play a role in gating. However, our MD simulations and electrophysiological data demonstrate that Tyr561 holds Glu_gate_ in the closed position and keeps intracellular protons from entering the pore. A limitation of McKiernan’s data is that only one subunit was analysed at 0 mV, under these conditions the coupling of fast (Glu_gate_) and common gates and the observation of gating transitions is limited because the open probability is almost null.

Is an electro-steric activation mechanism unique to CLC-2? We do not think so because voltage-dependent gating in CLC-0 and CLC-1 shows AMF behaviour, permeant anions facilitate gating, and non-permeant anions support voltage-dependent gating of CLC-1(Chen, 2003; Chen and Miller, 1996; Pusch et al., 1995; Rychkov et al., 2001, 1998). However, pore bifurcation could be key in fine-tuning activation. Protonation of Glu_gate_ in the bifurcated pores of CLC-1 and CLC-0 could be facilitated *after* Glu_gate_ interacts with the permeant anions. Thus, it is likely that a mechanism entailing electro-steric gating is present in all CLC Cl^−^ channels. In addition to CLC Cl^−^ channels, the K2P K^+^ channels that lack a voltage sensor domain also display voltage dependence due to the movement of K^+^ ions along the pore(Kopec et al., 2019; Schewe et al., 2016). Similarly, the voltage-dependent gating of viral K^+^ channel Kcv_NTS_ could rely on occupation by K^+^ of an external site(Rauh et al., 2018). Interestingly, the outward movement of ions could explain the activation of PIEZO channels by voltage alone(Moroni et al., 2018). Thus, coupling ion permeation to pore gating through electro-steric repulsion might be a more general activation mechanism than previously estimated(Hilgemann, 2020).

## Materials and Methods

### Homology model of mouse CLC-2 and preparation of membrane-protein ensembles

Initially, we constructed a model for the structure of the mouse CLC-2 Cl^−^ channel (908 aa) using the I-Tasser(Roy et al., 2010) server (https://zhanglab.ccmb.med.umich.edu/I-TASSER/) and the bovine CLC-K structure (5TQQ; Uniprot: E1B792; 687 aa) solved at 3.76 Å resolution using Cryo-EM(Park et al., 2017). This model structure was named CLC-2^CLC-K^. mCLC-2 is 45.7% identical to CLC-K within the transmembrane region. However, the pores of mCLC-2 and CLC-K differ in two residues; valine replaces Glu_gate_ in CLC-K. We built an additional model using as template the hCLC-1 structure determined by Cryo-EM (6COY; UniProt: P35523; 988 aa)(Park and MacKinnon, 2018) and the Modeller v9.19 software(Sali and Blundell, 1993). This model was named CLC-2^CLC-1^. The available hCLC-1 structure is based on 626 residues visible in the density maps. hCLC-1, a 988 residues long protein, was solved at 3.36 Å. CLC-2 and hCLC-1 are 56% identical within the transmembrane region and within the pore region, they differed by one amino acid. We measured the pore radius using CAVER 3.0(Chovancova et al., 2012).

The protonation state of ionisable residues at pH 7.3 was determined by PROPKA(Olsson et al., 2011). The protein, oriented using the Positioning of Proteins in Membrane server (PPM, http://opm.phar.umich.edu/server.php), was embedded in symmetric lipid bilayers of 1,2-dimyristoyl-sn-glycerol-3-phosphocholine (DMPC, CLC-2^CLC-K^) or 1-palmitoyl-2-oleoyl-sn-glycerol-3-phosphocholine (POPC, CLC-2^CLC-1^) generated using CHARMM-GUI(Jo et al., 2008) membrane builder (http://www.charmm-gui.org). The structures were solvated with water modelled by TIP3(Jorgensen et al., 1983) and 140 mM NaCl. Before MD simulation, the CLC-2^CLC-K^ and CLC-2^CLC-1^ systems consisted of 785 DMPC, 102741 TIP3, 800 Na^+^ and 416 Cl^−^ contained in a 170 × 170 × 158 Å^3^ simulation box, and of 554 POPC, 43283 TIP3, 110 Na^+^, and 130 Cl^−^ contained in a 176 × 171 × 100 Å^3^ box, respectively. Structures of Tyr561Phe and Tyr561Ala mutants were built on CLC-2^CLC-1^ using PyMol. Simulation boxes consisted of 554 POPC, 43271 TIP3, 111 Na^+^, and 131 Cl^−^ in a 170 × 167 × 102 Å^3^ box, and of 555 POPC, 43325 TIP3, 110 Na^+^, and 130 Cl^−^ in a 170 × 165 × 102 Å^3^ box for Tyr561Phe and Tyr561Ala, respectively.

### Molecular dynamics simulations

MD simulations were performed with the GPU-accelerated Gromacs 5.1 package(Berendsen et al., 1995) using the CHARMM36 force field(Klauda et al., 2010). The equations of motion were solved using the leapfrog algorithm with a time step of 2fs. The temperature was coupled to the Nosé−Hoover thermostat with a parameter τ_T_ = 0.2 ps at 300 K while the pressure was coupled to the Parrinello−Rahman barostat(Parrinello and Rahman, 1981) with a coupling parameter τ_P_ = 0.5 ps at 1 atm. Bond distances were kept rigid by using the LINCS algorithm. The electrostatic interactions were computed with the particle mesh Ewald (PME) approach(Darden et al., 1993) with a tolerance of 10^−6^ for the real space contribution, with a grid spacing of 1.2 Å, spline interpolation of order 4. In the isotropic NPT simulations, the real part of the Ewald summation and the LJ interactions were truncated at 12 Å and 10 Å, respectively. Long-range corrections for the LJ energy and pressure were included. All ensembles were minimized and equilibrated during 100 and 30 ns (WT and mutants), before implementing a potential difference across the membrane using the CompEl or the electric field protocols, respectively.

To perform CompEl we duplicated the CLC-2^CLC-K^ ensemble in the z-direction. Our CompEl simulation box consisted of three compartments separated by two DMPC bilayers embedding one CLC-2^CLC-K^ molecule each and solvated with 140 mM NaCl. Considering that charge (∆q), membrane capacitance (C_m_~1 µF/cm^2^), and membrane voltage (V) are related by the equation V=∆q/C_m_, we unbalanced the charge between compartments by moving Na^+^ ions to generate a voltage gradient across the membranes(Kutzner et al., 2011). A difference of 2 or 20 Na^+^ ions between central and side compartments is equivalent to apply +94 or +940 mV and −94 or −940 mV to the intracellular sides of ^+^CLC-2^CLC-K^ (channel 1 inserted in bilayer 1) and of ^−^CLC-2^CLC-K^ (channel 2 inserted in bilayer 2), respectively. This condition allowed us to analyse the CLC-2 closed state (positive voltages) and the open state (negative voltages) in the same MD simulation. Additionally, we applied a uniform electric field of −500 mV perpendicular to the membrane (+z direction is from intracellular to extracellular side), Ez. The last 200 ns of the MD simulation of WT CLC-2^CLC-K^ (Ez= V/L = −3.16 mV/Å, where V is the voltage and L is the size to the box on z-direction) and on Tyr561Phe or Tyr561Ala mutants (Ez = −4.9 mV/Å for both) was ran using this protocol. A protonated ^−^CLC-2^CLC-K^ molecule was inserted in a DMPC bilayer and solvated with 140 mM NaCl. Thus, our hybrid protocol for MD simulation consisted of 480 ns CompEl (144 ns at ±94mV followed by 340 ns at ±940 mV), protonation of ^−^CLC-2^CLC-K^ Glu_gate_, and of 200 ns applying an electric field of −500 mV to preserve the stability of the system. The protonation of Glu_gate_ (Glu213^H+^, Fig. 3) was carried out using CHARMM-GUI and equilibrated for 10 ns before applying an electrical field.

Gromacs tools, VMD and PyMol were used for analysis and visualization.

### Poisson-Boltzmann calculation

CLC-2^CLC-K^ in open and closed states was extracted at 565 ns from MD simulation. By solving the linearized Poisson-Boltzmann equation in CHARMM-GUI PBED solver(Im et al., 1998), the average electrostatic potential (EP) along the pore, identified by CAVER 3.0, was calculated on a 99 × 97 × 79 Å^3^ (1 Å grid spacing), with the CHARMM36 force field. The average potential was calculated on the transversal area determined by the pore radius. The dielectric constant for protein was 1, for membrane and membrane head group was 2, and for solvent containing 150 mM salt concentration was 80. The thickness of the membrane was 35 Å. *In silico* mutants were generated with CHARMM-GUI tools.

### Cell culture, transient expression and electrophysiological recordings

HEK-293 cells transfected with WT mouse ClC-2, mouse Tyr561Phe or mouse Tyr561Ala cDNAs were cultured and used for recording whole cell Cl^−^ currents (I_Cl_) as previously described(De Jesús-Pérez et al., 2016; De Santiago et al., 2005; Sánchez-Rodríguez et al., 2012, 2010). Control solutions used to record I_Cl_ contained (in mM): TEA-Cl 139, CaCl_2_ 0.5, HEPES 20 and D-mannitol 100 (external); and TEA-Cl 140, HEPES 20 and EGTA 20 (internal). HEPES was substituted by MES or phthalic acid to prepare solutions with low pH. The pH of these solutions was adjusted to 7.3 with TEA-OH. Average tonicity of external and internal solutions was 387.9±1.9 and 347.3±2.6 mosm/kg, respectively.

The mouse CLC-2 DNA inserted in the pGEM^®^-T vector was used for the heterologous expression in *Xenopus laevis* oocytes. The DNA was amplified, linearized with the Pme I enzyme (New England Biolabs, Inc., Ipswich, MA, USA) and transcribed in vitro using the T7 promoter mMESSAGE cRNA kit (Ambion, Austin, TX., USA). All mutations in this study were performed using standard PCR techniques and confirmed by sequencing the entire cassettes.

Procedures and methods involving *Xenopus laevis* frogs followed relevant guidelines and regulations as previously described(Rodríguez-Rangel et al., 2020). Briefly, oocytes contained in 1-3 mL of ovary lobes extracted via survival surgery were isolated by enzymatic digestion with collagenase type II (Worthington Biochemical Corp., NJ, USA) under mechanical agitation. After the isolation, each oocyte was injected with 40 ng of RNA encoding the WT mCLC-2 or mutants; oocytes were incubated for 2-7 days at 17 °C in a standard oocyte saline solution containing (in mM): 100 NaCl, 1 MgCl_2_, 10 HEPES, 2 KCl and 1.8 CaCl_2_ with 50 μg/mL gentamycin at pH 7.5. All chemical compounds utilized for this study were purchased from Sigma-Aldrich (Sigma-Aldrich Co., St. Louis, MO, USA).

I_Cl_ from HEK cells was recorded using a protocol that consisted of holding the cell at 0 mV, then applying a voltage between +60 or +200 to −200 mV in 20 mV steps and then returning the cell to +60 or +80 mV. All experiments were performed at room temperature (21-23°C). To determine the effect of AMF on ClC-2 activation, we used internal solutions containing 0.0, 0.25, 0.5, 0.75, and 1.0 acetate mole fractions. I_Cl_ was recorded using pClamp 10 and sampling at 500 kHz. Otherwise, currents were sampled using pClamp V8. To avoid electrode polarization while using internal solutions with low Cl^−^ we used an electrode embedded in a 3 M KCl/3% agar jacket(Shao and Feldman, 2007). Membrane and reversal potentials were corrected off-line using liquid junction potentials experimentally measured(Neher, 1992).

Electrophysiology measurements in *Xenopus laevis* oocytes were conducted using the cut-open oocyte voltage clamp methodology(Siefani and Bezanilla, 1998). The internal recording solution contained (in mM): 136 NMDG (N-methyl-D-glucamine)-HCl, 2 MgCl_2_, 10 EGTA, 10 HEPES. The external recording solution was composed (in mM) by 130 NMDG-HCl, 4 MgCl_2_, 1 BaCl_2_ and 10 HEPES. Both solutions were adjusted to pH 7.3 with 1M NMDG. I_Cl_ was amplified and digitized using the Oocyte Clamp Amplifier CA-1A (Dagan Corporation, Minneapolis, MN, USA) and the USB-1604-HS-2AO Multifunction Card (Measurement Computing, Norton, MA, USA). The acquisition system was controlled by the GpatchMC64 program (Department of Anesthesiology, UCLA, Los Angeles, CA, USA) via a personal computer. Recordings sampled at 100 kHz and filtered at 10 kHz were obtained at room temperature (21–23 °C).

### Analysis

The voltage dependence of the open probability (P_A_) was determined by constructing curves of P_A_ (=G/G_max_) versus voltage (P_A_(V)). Conductance (G) at each V was calculated as I_Cl_/(V-V_r_), where V is the membrane potential and V_r_ is the reversal potential. The maximum conductance (G_max_) was estimated before normalization by fitting the G against V curves with the Boltzmann equation:

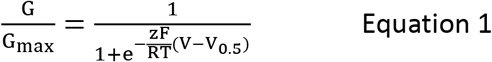

where z is the apparent charge, F is the Faraday constant, R is the gas constant, T is the temperature and V_0.5_ is the V needed to reach P_A_ = 1/2. Instantaneous current-voltage plots were constructed using the magnitude of tail currents recorded at different V and fitted to a linear function to determine V_r_.

The voltage-dependence of the open probability of Glu_gate_ (P_GluGate_)and common (P_C_) gates was calculated as described before(De Santiago et al., 2005). Briefly, we used a 15 ms hyperpolarization pulse to −200 mV to completely open Glu_gate_ (time constant ≤3ms) so P_Glugate_ ~1. When I_Cl_ reaches steady-state P_A_ = P_Glugate_ * P_C_, before the hyperpolarization step to −200 mV I_Cl_ obeys the relationship I_o_ = 2NiP_Glugate_*P_C_, and after the hyperpolarization step I_Cl_ is given by I_ahs_ ≈ 2NiP_C_. Here, N is the total number of channels, i is the single-channel current, all multiplied by 2 because is a two-pore channel. Hence, P_Glugate_ can be calculated as I_o_/I_ahs_ and P_C_ = P_A_/P_Glugate_. Cut open voltage-clamp recordings were included in the analysis after determining that the reversal potential values of I_Cl_ were near the predicted Nernst potential for Cl^−^. I_Cl_ recordings were analysed with the software Analysis (Department of Anesthesiology, UCLA, Los Angeles, CA, USA).

Figures and fits were carried out using Origin (Origin Lab, Northampton, MA). Experimental data are plotted as mean ± SEM of n (number of independent experiments). Dashed black lines in each Figure indicate I_Cl_ = 0. Where necessary, a paired Student t-test was used to evaluate significant differences at P<0.05 between data sets.

## ACKNOWLEDGEMENTS

The authors thank Dr Patricia Perez-Cornejo for critical comments, Carmen Y. Hernandez-Carballo for technical assistance. The work was supported by grants 219949 and FC-2016-01-1955 from CONACyT, Mexico. RG-G is a recipient of a Graduate Student Fellowship #726278 from CONACyT, Mexico. JJDeJ-P was supported by a Graduate Student Fellowship and Postdoctoral Fellowship 234820 and 711128 from CONACyT, Mexico, respectively.

## AUTHOR CONTRIBUTIONS

JJDeJP Designed research, performed patch-clamp experiments, performed MD simulations, analysed data and wrote the manuscript.

VDRJ Performed patch-clamp experiments, analysed data and review the manuscript.

ILGH Performed patch-clamp experiments

GAMM Performed MD simulations and analysed the data

JESR, RG-G Performed electrophysiological experiments in X. oocytes, analysed the data and wrote the manuscript.

JA Designed research, analysed data, wrote the paper, and secure funds.

