## Supplemental Data for "Electro-Steric Mechanism of CLC-2 Chloride Channel Activation"

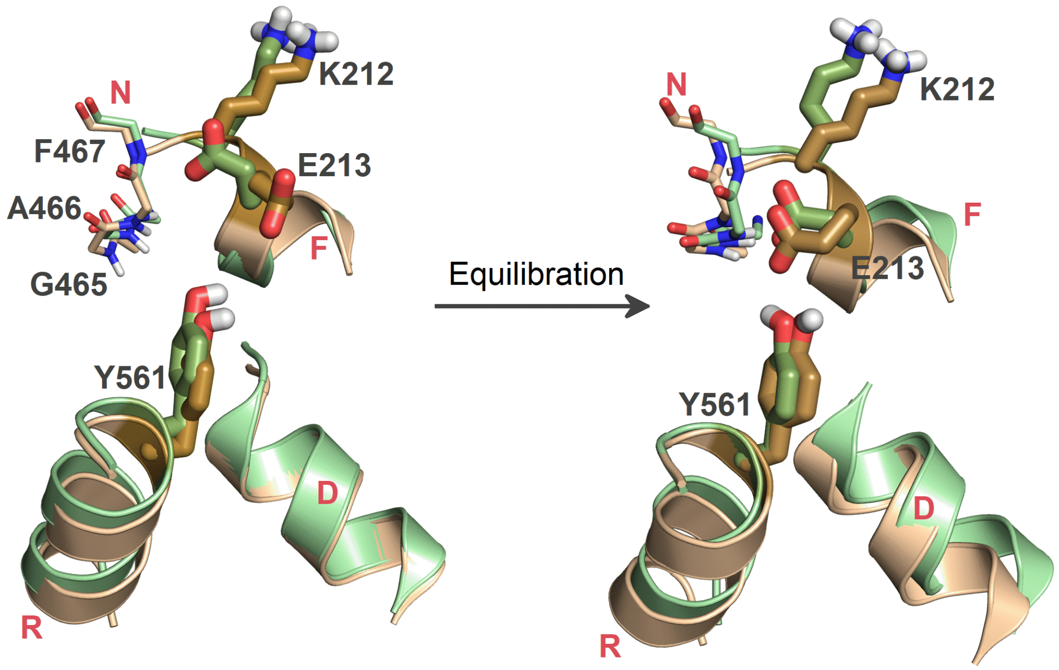


**Supplementary Fig. 1 The homology structure of the canonical pores of CLC-2$^{CLC-K}$ (green) and CLC-2$^{CLC-1}$ (orange) is the same.**

The model structures CLC-2$^{CLC-K}$ (generated with I-Tasser using the bovine CLC-K structure 5TQQ) and CLC-2$^{CLC-1}$ (generated with Modeller using hCLC-1 structure 6COY) of the mouse CLC-2 $Cl^-$ channel are superimposed before (on the left) or after equilibration (on the right). Before equilibration, CLC-2$^{CLC-K}$ and CLC-2$^{CLC-1}$ were embedded in a DMPC and POPC bilayer, hydrated with 140 mM NaCl, and used for MD simulation at 0 mV during 100 ns and 30 ns, respectively. Lys212, Glu213, and Tyr561 are represented as thick sticks and the $_{465}$GAF$_{467}$ loop as thin sticks.

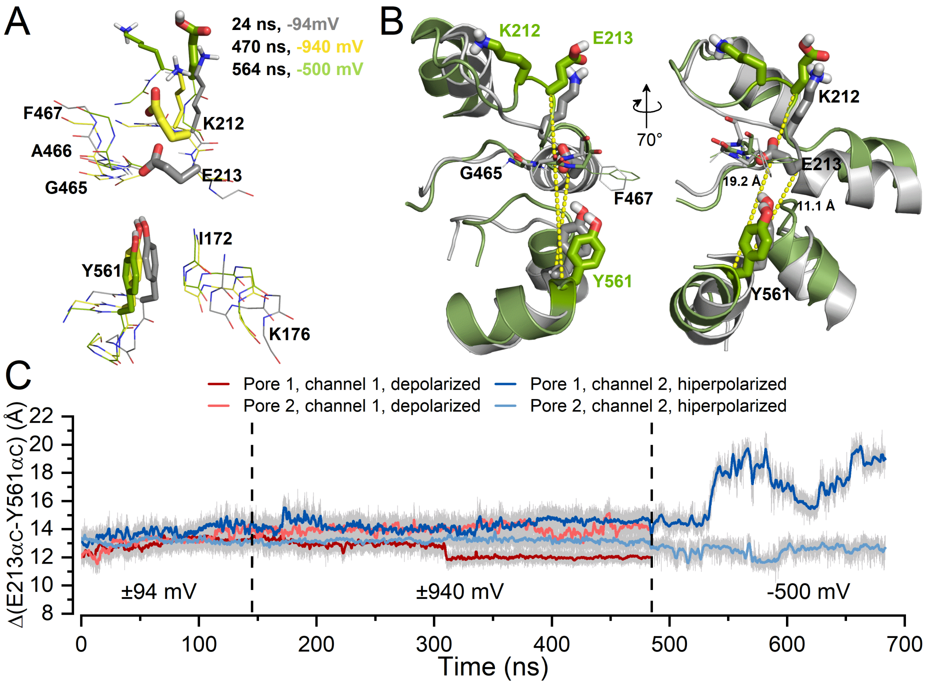


**Supplementary Fig. 2 Conformational changes underwent by residues lining the canonical pore.**

A. Snapshots to show the conformations adopted by $Glu_{gate}$ (Glu213) in pore 1 of $^{-}$CLC-$2^{CLC-K}$ in the presence of $Cl^-$ at the indicated simulation times and transmembrane voltages.

Grey = 24 ns and -94mV

Yellow = 470 ns, -940 mV

Lemon green = 564 ns, at -500 mV.

$Glu_{gate}$ (E213) and Tyr561 are represented as thick sticks and Lys212 as thin sticks.

B. Closed (grey) and open states (green) of the CLC-$2^{CLC-K}$ pore are compared after 100 ns of equilibration and 564 ns into the MD simulation. On the right side, the pore was rotated

70°. Yellow dash lines indicate the distances from the α-carbon of Tyr561 to the α-carbon of $Glu_{gate}$. The chloride ion is not shown for clarity.

C. The distance between Tyr561 and $Glu_{gate}$ increased as $Cl^-$ exited the pore. The distance from the α−carbon of Tyr561 to the α−carbon of Glu213 (yellow dash lines in B) as a function of time. The distance was measured for both pores at -94, -940, and -500 mV (blue traces) and at +94, and +940 mV (red traces).

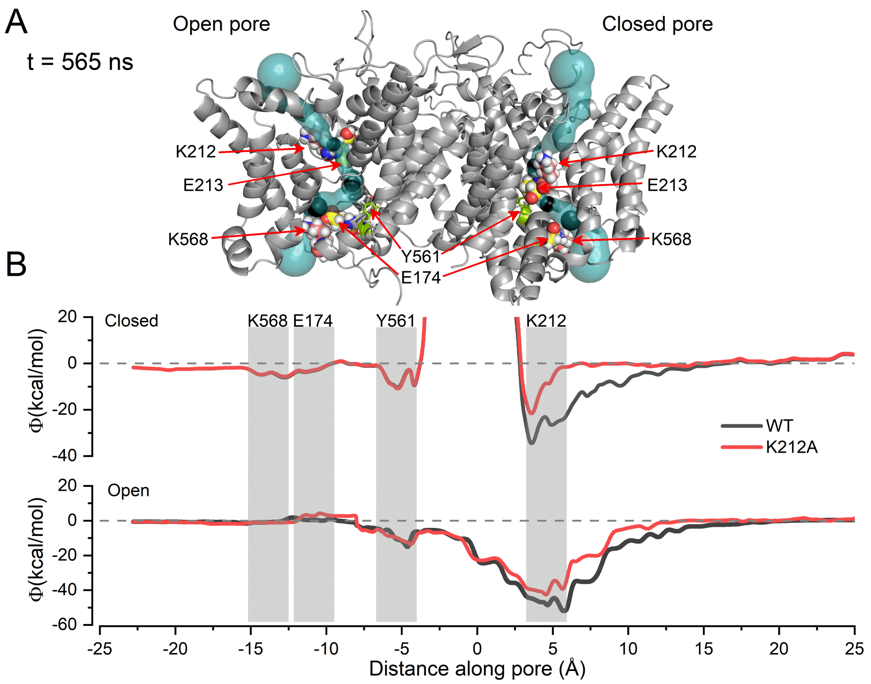


**Supplementary Fig. 3 Electrostatic potential along the permeation pathway in the closed and open states.**

A. $^{-}$CLC-2$^{CLC\text{-}K}$ protein at 565 ns. The permeation pathways (blue surfaces) were determined by CAVER software in open (left) and closed (right) pores. Glu174 and Glu213 (yellow spheres), Lys212 and K568 (pink), and Tyr561 (lemon green) are shown in space-filling representation.

B. The electrostatic potentials along the closed (upper plot) and open (lower plot) pores of WT CLC-2 (grey) and Lys212Ala (red) mutant channels were calculated using the linearized Poisson-Boltzmann equation and the trajectories determined by CAVER in A. Rectangle shadows limit the corresponding region of Lys568, Glu174, Tyr561, and Lys212 residues highlighted in A. The electrostatic potential at Lys2122 in pore 2 (closed) and pore 1 (open) of $^{-}$CLC-2$^{CLC\text{-}K}$, were -26 Kcal/mol and -52 Kcal/mol, respectively.

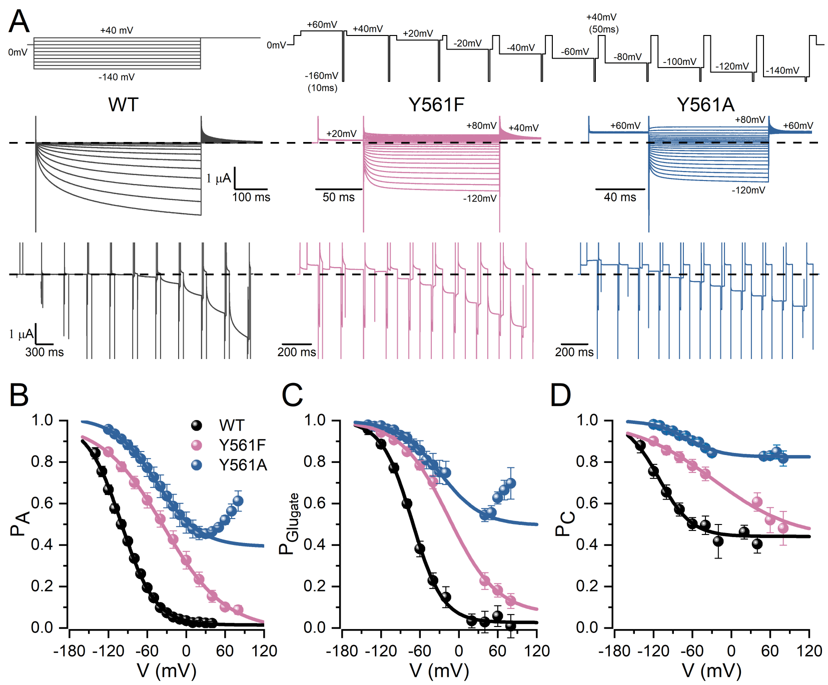


**Supplementary Fig. 4 Functional characterization of WT, Tyr561Phe-CLC-2 and Tyr561Ala-CLC-2 chloride channels using the cut-open oocyte voltage clamp methodology.**

A. Representative chloride recordings (color-coded) acquired from three different oocytes expressing WT CLC-2, Tyr561Phe-CLC-2 and Tyr561Ala-CLC-2. Chloride currents were first elicited by hyperpolarization steps from 40 (or 80) to -140 mV (or -120) in 10 mV increments followed by a single depolarized pulse at 40 (or 60) mV to record the tail currents (upper left). Just after, a strong hyperpolarization to -160 mV and 10 ms duration was delivered as interpulse between pulses that varied the membrane voltage between 60 to -140 mV. The inter-pulse force $P_{Glugate} \approx 1$ to calculate $P_{Glugate}$ and $P_C$ as described in Methods. Currents were assessed at pHi = pHo = 7.3 and [Cl-]i = [Cl-]o = 140 mM. Dash line indicates $I_{Cl} = 0$.

B. Apparent open probability ($P_A$) *vs* voltage.

C. Open probability of Glugate ($P_{Glugate}$) *vs* voltage.

D. Open probability of common gate ($P_C$) *vs* voltage.

Open probabilities of WT CLC-2, Tyr561Phe-CLC-2, and Tyr561Ala-CLC-2 (colour-coded) were determined as described in the methods section. Continuous lines represent fits with the Boltzmann equation to determine the voltage-dependent parameters ($V_{0.5}$ and z) listed in Table 1.

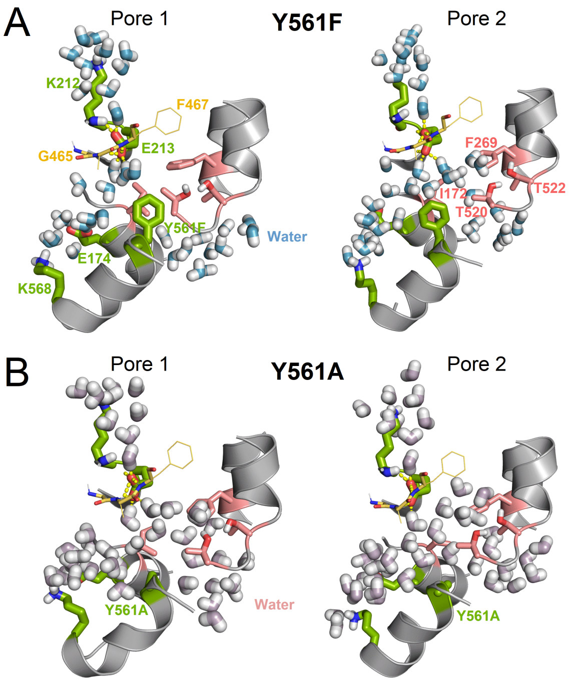

**Supplementary Fig. 5 Mutating Tyr561 to Phe or Ala disarrange the Tyr561-$H_2O$-$Glu_{gate}$ gate in both pores of $^-$CLC-2$^{CLC-K}$.**

Phe561 (in A), Ala561 (in B), and $Glu_{gate}$, Ly568, Glu174, and Lys212 (in A and B) are shown in green whereas residues forming the hydrophobic gate are depicted in pink. The Tyr561Ala mutation causes the pore to become flooded at the branch connecting the canonical pore with the alternative pathway.
